# Engineered immunomodulatory extracellular vesicles derived from epithelial cells acquire capacity for positive and negative T cell co-stimulation in cancer and autoimmunity

**DOI:** 10.1101/2023.11.02.565371

**Authors:** Fernanda G. Kugeratski, Valerie S. LeBleu, Dara P. Dowlatshahi, Hikaru Sugimoto, Kent A. Arian, Yibo Fan, Li Huang, Danielle Wells, Sergio Lilla, Kelly Hodge, Sara Zanivan, Kathleen M. McAndrews, Raghu Kalluri

## Abstract

Extracellular vesicles (EVs) are generated by all cells and systemic administration of allogenic EVs derived from epithelial and mesenchymal cells have been shown to be safe, despite carrying an array of functional molecules, including thousands of proteins. To address whether epithelial cells derived EVs can be modified to acquire the capacity to induce immune response, we engineered 293T EVs to harbor the immunomodulatory CD80, OX40L and PD-L1 molecules. We demonstrated abundant levels of these proteins on the engineered cells and EVs. Functionally, the engineered EVs efficiently elicit positive and negative co-stimulation in human and murine T cells. In the setting of cancer and auto-immune hepatitis, the engineered EVs modulate T cell functions and alter disease progression. Moreover, OX40L EVs provide additional benefit to anti-CTLA-4 treatment in melanoma-bearing mice. Our work provides evidence that epithelial cell derived EVs can be engineered to induce immune responses with translational potential to modulate T cell functions in distinct pathological settings.

## Introduction

Extracellular vesicles (EVs) are membrane-enclosed entities released by cells into neighbor tissues and body fluids ^1^. EVs carry an array of functional molecules, which include nucleic acids ^2^, proteins ^3^, post-translational modifications ^4^, amino acids ^5^, lipids ^6^, and metabolites ^7^. Notably, EVs can function as signaling conduits between cells and across tissues, through the transfer of their molecular constituents, or by initiating receptor-mediated signaling in the target cells ^8–13^.

EVs have emerged as regulators of immune cell functions in physiology and in pathologies where the immune system plays a fundamental role, such as cancer, auto-immune diseases, and sepsis ^14–16^. From a translational standpoint, EVs can be modified to carry biomolecules and agents that modulate the immune system to promote anti-tumor immunity ^17^. In fact, EVs harboring immunomodulatory proteins and cytokines ^18–21^, chimeric antigen receptor ^22^, alarmin ^23^, and STING agonist ^24^ have shown promising anti-tumor activity in preclinical models of cancer. Additionally, immunomodulatory proteins were previously detected in patient derived EVs, linked to immunoregulation, and proposed to have a prognostic value in cancer ^9,14,25^. However, a comprehensive functional characterization of EV-mediated engagement of positive and negative T cell co-stimulation, as well as their translational potential have been underexplored.

Studies employing cross-species, dendritic cell, allogenic mesenchymal stem cells (MSC), or embryonic kidney epithelial cells (293T cells)-derived EVs, suggest that despite possessing thousands of proteins including MHC class I, EVs were well tolerated, did not elicit toxicity nor adverse immune responses ^26–29^. We hypothesized that the concentration of any individual protein in a single EV might be too low to elicit an immune response. To test this hypothesis, we engineered EVs to harbor T cell immunomodulatory proteins, namely CD80, OX40L and PD-L1. We validated the feasibility of our approach to generate EVs harboring high levels of the immunomodulatory proteins of interest with capacity to launch T cell-dependent adaptive immune response. We established the functionality of the engineered EVs to engage positive and negative T cell co-stimulation in both human and murine T cells. Notably, in the setting of cancer and auto-immune hepatitis, the engineered EVs modulate T cell functions and alter disease progression. Our findings provide evidence that epithelial EVs can be engineered to harbor immunomodulatory proteins that engage positive and negative signals in T cells and have the potential to be explored therapeutically in the setting of cancer and auto-immune diseases.

## Results

### EVs can be engineered to harbor immunomodulatory proteins

To establish the role of EVs in engaging positive and negative T cell co-stimulation, we engineered human 293T cell-derived EVs to harbor the CD80, OX40L and PD-L1 proteins. Owing the function of these immune checkpoint proteins in T cell biology, we hypothesized that CD80-containing EVs may display a dual role in T cell functions, whereby upon binding to CD28 receptor, EV-resident CD80 would activate T cells, and upon binding to CTLA-4, it would suppress T effector functions. In addition, we hypothesized that EVs harboring OX40L would functionally activate the OX40 pathway in T cells to enable T cell activation; and PD-L1-containing EVs, would inhibit T cell functions upon engagement of the PD-1 pathway (**Fig. 1a**). Based on these premises, we generated HEK293T cells overexpressing the human CD80, human OX40L and human PD-L1 proteins, hereafter referred to as hCD80, hOX40L, and hPD-L1. We validated the overexpression of hCD80, hOX40L, and hPD-L1 in comparison to WT cells at the transcript level by RT-qPCR, and at the protein level by western blot and flow cytometry (**Fig. 1, b-d**). Notably, we observed that the EVs released by hCD80, hOX40L, and hPD-L1-overexpressing cells harbor high levels of the hCD80, hOX40L, and hPD-L1 proteins, respectively, as opposed to their WT counterpart, as demonstrated by western blot, and flow cytometry-based analysis of non-permeabilized EVs bound to beads (**Fig. 1c, d**). Together, these results demonstrate that EVs can be successfully engineered to carry immunomodulatory checkpoint proteins capable of eliciting a T cell-dependent adaptive response.

**Fig. 1.**
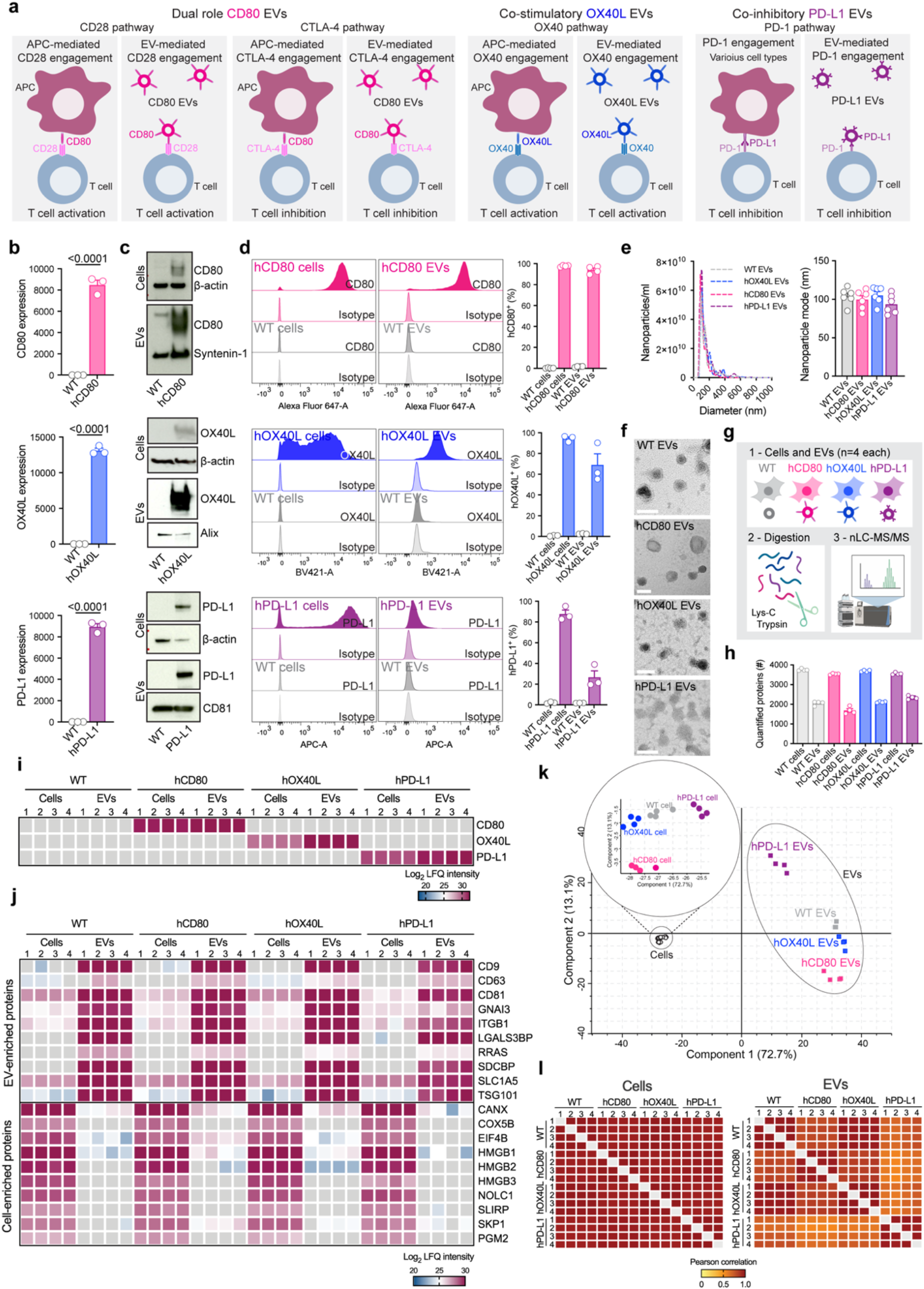
EVs can be modified to harbor high levels of the immunomodulatory checkpoint proteins CD80, OX40L and PD-L1. (**a**) Schematic representation of APC/cell-mediated and of EV-mediated engagement of the CD28/CTLA-4, OX40 and PD-1 pathways in T cells. (**b**) Relative expression of human *CD80*, *OX40L* and *CD274* (PD-L1), in parental cells determined by RT-qPCR. Bar graph shows mean +/- s.e.m. of fold-change from n=3 biological replicates normalized to *GAPDH.* Statistical significance was determined by two-tailed unpaired t-test, and p-values are shown. (**c**) Human CD80, OX40L and PD-L1 and loading control proteins in parental cells and EVs probed by western blot. (**d**) Flow cytometry-based evaluation of overexpressing proteins at the surface of parental cells and EVs. Overlaid histograms for each protein show the profile for WT and engineered cells and EVs stained with isotype control and with the antibody of interest. The accompanying bar graphs show mean +/- s.e.m. of percentage of positive cells and of positive beads (for EV analyses). Individual data points from n=3-4 biological replicates are shown. (**e**) Representative NTA profile of WT and engineered EVs. Bar graph shows mean +/- s.e.m. of nanoparticle mode measurements from n=6 biological replicates. (**f**) Morphology of WT and engineered EVs determined by TEM. Scale bar = 100 nm. (**g**) Schematic representation of the workflow utilized for the MS-based proteomics analysis of parental cells and EVs. (**h**) Bar graph shows mean +/- s.e.m. of number of proteins quantified in cells and EVs by MS, results from n=4 biological replicates. (**i**) Heatmap of CD80, OX40L and PD-L1 protein levels in parental cells and EVs determined by MS, results from n=4 biological replicates. Gray colored squares represent no detection. (**j**) Heatmap of EV markers and exclusion markers in parental cells and EVs determined by MS, results from n=4 biological replicates. Gray colored squares represent no detection. (**k**) PCA analysis of proteomes from parental cells and EVs, results from n=4 biological replicates. (**l**) Heatmaps show Pearson correlation of proteomes from parental cells (left) and EVs (right), results from n=4 biological replicates.

To investigate whether the overexpression of hCD80, hOX40L, and hPD-L1 influences biophysical characteristics of EVs, we evaluated the size distribution and the morphology of the nanoparticles released by hCD80, hOX40L, and hPD-L1-overexpressing cells using nanoparticle tracking analysis (NTA) and transmission electron microscopy (TEM). We observed that the size distribution, the nanoparticle mode, and the morphology of hCD80, hOX40L, and hPD-L1 EVs was comparable to WT EVs (**Fig. 1e, f**). Further, to establish whether the overexpression of hCD80, hOX40L, and hPD-L1 alters the molecular composition of cells and/or their shed EVs, we used an unbiased and global mass spectrometry (MS)-based proteomics approach based on label-free quantification (**Fig. 1g**). The total number of proteins quantified in the proteome of cells and EVs was similar across WT, hCD80, hOX40L, and hPD-L1, with ∼4,000 and ∼2,000 proteins quantified in cells and EVs, respectively (**Fig. 1h**). Taken together, these findings suggest that the overexpression of the immunomodulatory proteins hCD80, hOX40L, and hPD-L1 did not induce major changes in the size, morphology, and in the number of proteins detected in parental cells and EVs.

Global proteomic analysis revealed a specific enrichment of hCD80, hOX40L, and hPD-L1 proteins in the engineered cells and EVs in comparison to the WT counterparts, thus validating our previous observations from western blot-based and flow cytometry-based analyses (**Fig. 1i**). Additionally, we exploited the comprehensive proteomic dataset generated here to evaluate the presence and abundance of EV markers and exclusion markers ^3^. This analysis revealed that several exclusion markers (CANX, COX5B, EIF4B, HMGB1, HMGB2, HMGB3, NOLC1, SLIRP, SKP1, and PGM2) were abundant in the parental cells, and low abundant or undetected in EVs released by these cells (**Fig. 1j**). On the other hand, the EV markers SDCBP (Syntenin-1), TSG101, CD9, CD63, CD81, SLC1A5, GNAI3, ITGB1, LGALS3BP and RRAS were highly abundant in EVs as opposed to the parental cells (**Fig. 1k**).

To gain additional insights into the proteomic composition of hCD80, hOX40L, and hPD-L1 cells and EVs in relation to their WT counterpart, we employed principal component analysis (PCA), Pearson correlation and hierarchical clustering analyses. Together, these analyses demonstrated that the proteomic composition of hCD80, hOX40L, and hPD-L1 cells was comparable to the profile of WT cells (**Fig. 1k-l, Extended Data Fig. 1a**). Additionally, our analyses suggest that the proteomic composition of hCD80 and hOX40L EVs was closely related to the WT EVs, whereas hPD-L1 EV samples cluster separated and had lower correlation to the other EVs evaluated (**Fig. 1k-l, Extended Data Fig. 1a**). This suggests that high levels of PD-L1 in EVs may influence the packaging of other proteins in EVs.

Further, we also generated HEK293T cells that produce EVs harboring the murine CD80, murine OX40L and murine PD-L1 proteins, hereafter referred to as mCD80, mOX40L, and mPD-L1. The transcript of mCD80, mOX40L, and mPD-L1 was detected at high levels in the overexpressing cells, but not in the WT counterparts (**Extended Data Fig. 1b**). Moreover, we validated that the engineered parental cells and their EVs display the immunomodulatory checkpoints at their surface by flow cytometry (**Extended Data Fig. 1c**), and that the engineering process did not induce major biophysical changes in terms of size distribution and morphology of mCD80, mOX40L, and mPD-L1 EVs (**Extended Data Fig. 1d, e**).

To gain insights into the predicted specificity of the signaling elicited by the engineered EVs in T cells, we sought to understand whether the human 293T parental cells and EVs contain other co-stimulatory (OX40L, iCOSL, 4-1BBL, CD70), co-inhibitory (Gal-9, HVEM, PD-L1, PD-L2) and dual-role (CD80) immune checkpoint proteins. Using flow cytometry-based analysis of parental cells and EVs, we observed that high levels of human CD80, OX40L, and PD-L1 proteins were readily detected in the engineered hCD80, hOX40L, and hPD-L1 cells and EVs, respectively, but not in the WT and in the mCD80, mOX40L, and mPD-L1 engineered cells and EVs (**Extended Data Fig. 1g, h**). Moreover, low levels of CD70, Gal-9, HVEM, iCOSL, PD-L1 and 4-1BBL were detected on the surface of cells (**Extended Data Fig. 1g**). From all the immune checkpoints evaluated, the co-stimulatory protein iCOSL was detected at low levels in the EVs (**Extended Data Fig. 1h**). We also conducted the flow cytometry-based evaluation of the putative EV markers CD9, CD63 and CD81 in the WT, hCD80, hOX40L, hPD-L1, mCD80, mOX40L, and mPD-L1 EVs. We observed that all the EVs evaluated contain high levels of these tetraspanins (**Extended Data Fig. 1f**). Taken together, our results indicate that the overexpressing proteins of interest are the prevalent immunomodulatory signal in the engineered EVs.

### Engineered EVs elicit positive and negative co-stimulation in human T cells

To determine whether the engineered EVs can functionally engage their cognate pathways in T cells, we performed *ex vivo* assays using human T cells derived from PBMCs. As a first step to characterize our *ex vivo* system, we profiled the kinetics of the receptors of interest, namely CD28, CTLA-4, OX40 and PD-1 in human T cells every 24 hours, from 0 to 168 hours post T cell activation. For the evaluation of CD28 and CTLA-4 surface levels over time, T cells were stimulated with anti-CD3e, which mimics antigen presentation. For the evaluation of OX40 and PD-1 surface protein over time, in addition to CD3e, T cells were concomitantly stimulated with anti-CD28 (**Extended Data Fig. 2a**). Using flow cytometry, we observed that in culture both human CD4^+^ and CD8^+^ T cells were positive at their surface for CD28 in all time points evaluated, but not for CTLA-4, suggesting that under these conditions, the engineered hCD80 EVs potentially elicit positive T cell co-stimulation, considering the high availability of surface CD28 receptor as opposed to CTLA-4. As for the OX40 receptor, upon activation, the surface OX40 increased, peaked at 24-48 hours, and dramatically decreased after 96 hours. PD-1 receptor on the T cell surface increased upon activation of both CD4^+^ and CD8^+^ T cells over time and peaked at the later time point of 168 hours (**Extended Data Fig. 2c**). The insights gained with the kinetics of the CD28, CTLA-4, OX40 and PD-1 receptors in human T cells allowed us to design *ex vivo* assays leveraging the availability of these receptors to test the functionality of the engineered EVs in modulating CD4^+^ and CD8^+^ T cell functions (**Fig. 2a**).

**Fig. 2.**
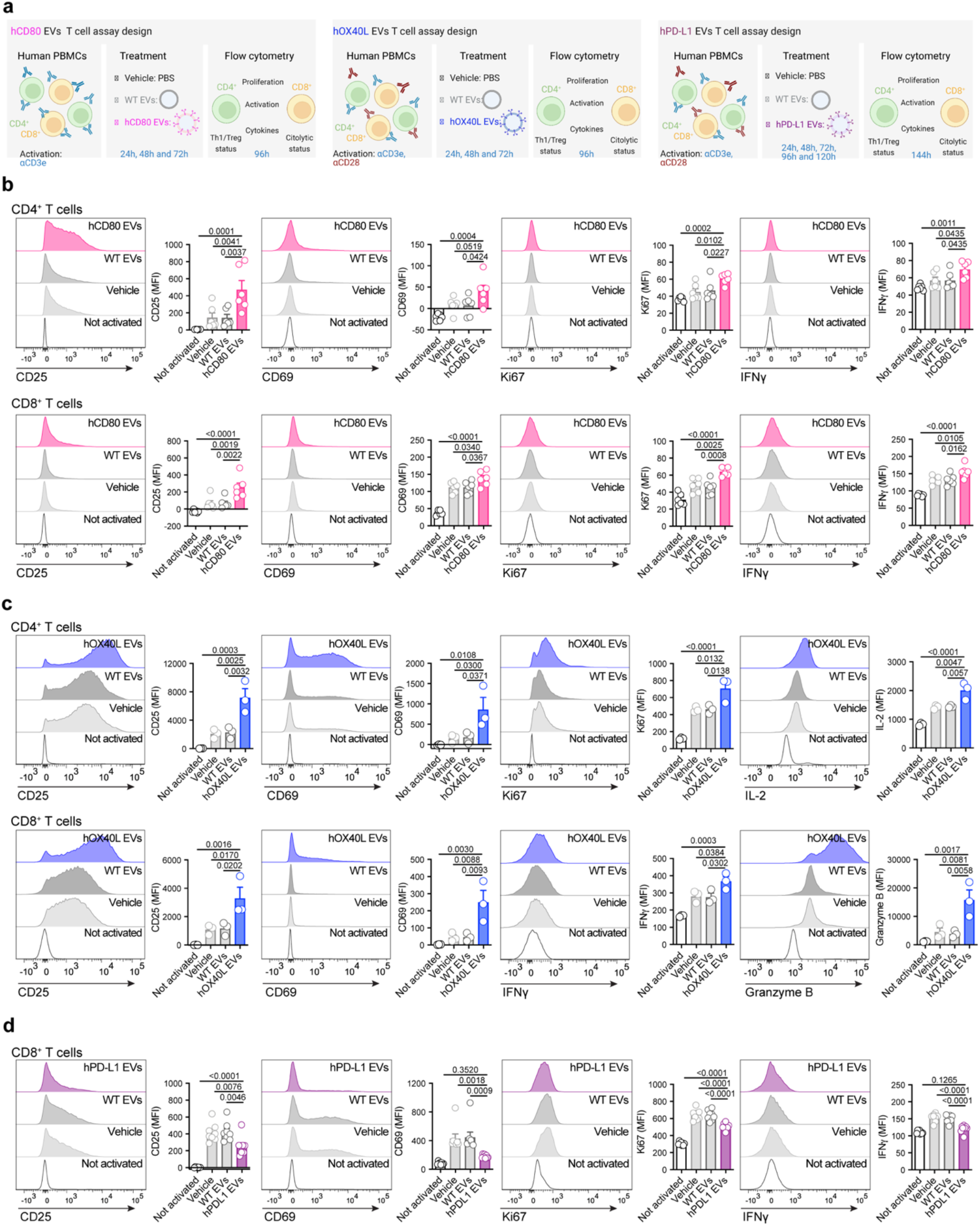
Engineered EVs elicit positive and negative co-stimulation in human T cells. (**a**) Schematic representation of the experimental design deployed to investigate the functionality of engineered EVs ex vivo using human PBMCs. (**b**) Representative histograms of activation markers (CD25 and CD69), proliferation marker (Ki67), and IFNγ in human CD4^+^ T cells and CD8^+^ T cells treated with hCD80 EVs. Accompanying bar graphs show mean +/- s.e.m. of MFI from n=6 PBMC samples. (**c**) Representative histograms of activation markers (CD25 and CD69), proliferation marker (Ki67), and IL-2 in human CD4^+^ T cells, and activation markers (CD25 and CD69), IFNγ, and cytolytic marker (Granzyme B), in human CD8^+^ T cells treated with hOX40L EVs. Accompanying bar graphs show mean +/- s.e.m. of MFI from n=3 PBMC samples. (**d**) Representative histograms of activation markers (CD25 and CD69), proliferation marker (Ki67), and IFNγ in human CD8^+^ T cells treated with hPD-L1 EVs. Accompanying bar graphs show mean +/- s.e.m. of MFI from n=8 PBMC samples. Statistical significance was determined using ordinary one-way ANOVA, p-values are shown. Statistical significance defined as p < 0.05.

We treated human T cells with vehicle (PBS), WT EVs, or engineered EVs (hCD80, hOX40L, hPD-L1) and evaluated several parameters for T cell functions, namely activation (CD25 and CD69), proliferation (Ki67), cytokines (IL-2 and IFNγ), and cytolytic potential (Granzyme B). Both CD4^+^ and CD8^+^ T cells responded to hCD80 EVs by increasing the CD25, CD69, Ki67 and IFNγ (**Fig. 2b**). Human CD4^+^ T cells treated with hOX40L EVs displayed a significant increase in CD25, CD69, Ki67 and IL-2, whereas CD8^+^ T cells treated with hOX40L EVs significantly increased CD25, CD69, IFNγ, and Granzyme B (**Fig. 2c**). As for PD-L1-containing EVs, we observed that human CD8^+^ T cells significantly decreased their activation, proliferation and cytokine production when treated with hPD-L1 EVs in comparison to controls (**Fig. 2d**), as opposed to CD4^+^ T cells, in which no major changes in functional markers were observed (**Extended Data Fig. 3b**).

Taken together, our results demonstrate that EVs harboring immune checkpoint proteins can functionally elicit positive and negative co-stimulation of human T cells *ex vivo*.

### Engineered EVs elicit positive and negative co-stimulation in murine T cells

Next, to understand whether the engineered EVs can also functionally engage their cognate pathways in murine-derived T cells, we performed *ex vivo* assays using T cells isolated from mouse spleen. To characterize the murine *ex vivo* system, we measured the kinetics of CD28, CTLA-4, OX40 and PD-1 in murine T cells post activation (**Extended Data Fig. 2b**). Both murine CD4^+^ and CD8^+^ T cells were positive for CD28 from 24 hours to 168 hours post activation, with a peak observed at 48 hours. Surface CTLA-4 was low in both CD4^+^ and CD8^+^ T cells for all time points evaluated. This suggests that under these conditions, the engineered mCD80 EVs would likely trigger positive T cell stimulation, considering the high availability of surface CD28 receptor in T cells in comparison to CTLA-4. The OX40 receptor increased upon murine T cell activation, peaked at 48 hours, and dramatically decreased after 72 hours. The surface PD-1 receptor increased upon activation of both CD4^+^ and CD8^+^ T cells and peaked at 48 hours post activation (**Extended Data Fig. 2d**). Since the murine CD28, OX40 and PD-1 receptors were positive in T cells at 24 hours, and peaked at 48 hours post-activation, for our murine *ex vivo* assays, T cells were treated with the engineered EVs at 24 hours and 48 hours post-activation, and the functional markers evaluated at 72 hours by flow cytometry (**Fig. 3a**).

**Fig. 3.**
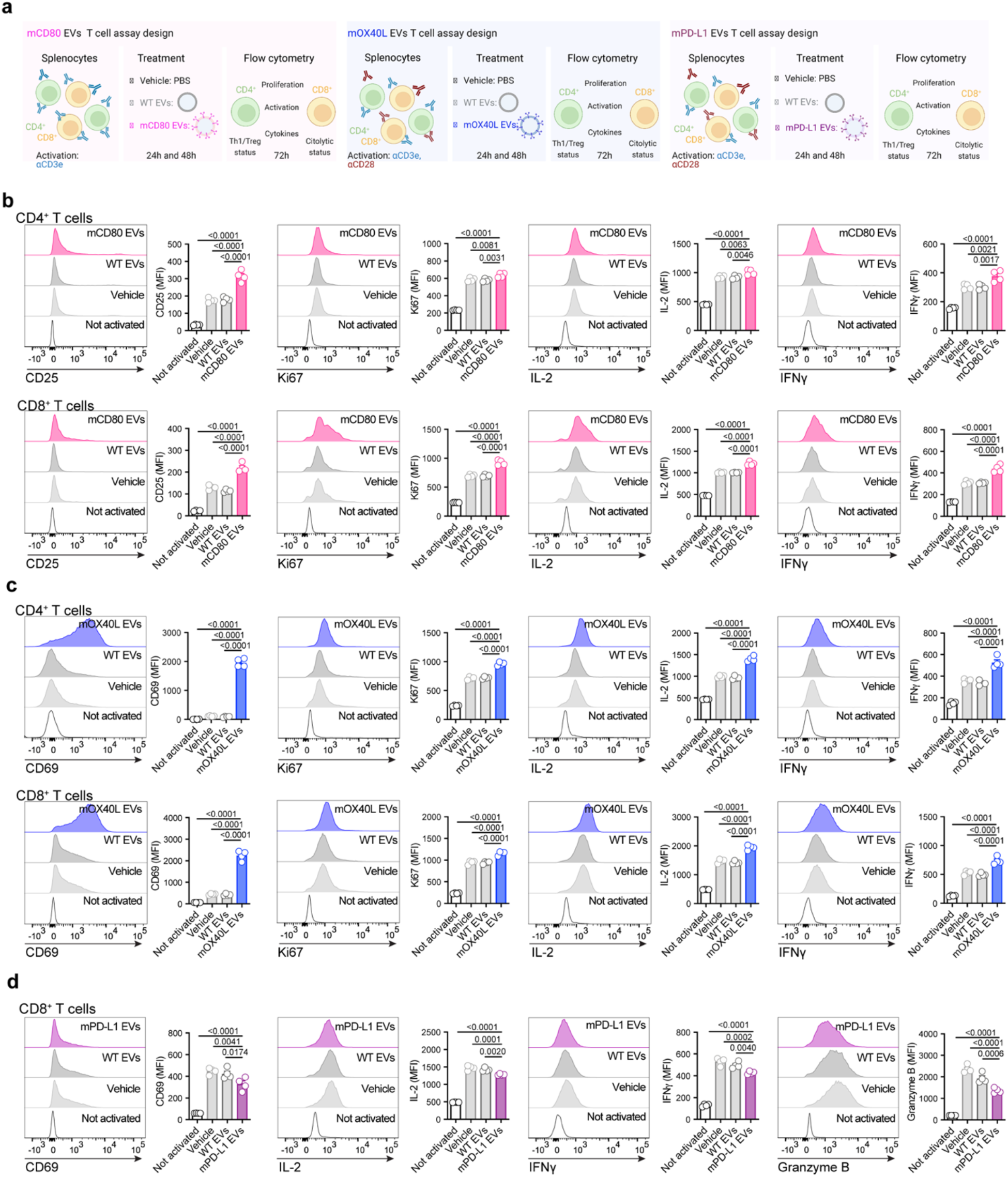
Engineered EVs elicit positive and negative co-stimulation in murine T cells. (**a**) Schematic representation of the experimental design used to investigate the functionality of engineered EVs ex vivo using murine splenocytes. (**b**) Representative histograms of activation marker (CD25), proliferation marker (Ki67), and cytokines (IL-2 and IFNγ) in murine CD4^+^ T cells and CD8^+^ T cells treated with mCD80 EVs. Accompanying bar graphs show mean +/- s.e.m. of MFI from n=4 mice. (**c**) Representative histograms of activation marker (CD69), proliferation marker (Ki67), and cytokines (IL-2 and IFNγ) in murine CD4^+^ T cells and CD8^+^ T cells treated with mOX40L EVs. Accompanying bar graphs show mean +/- s.e.m. of MFI from n=4 mice. (**d**) Representative histograms of activation marker (CD69), cytokines (IL-2 and IFNγ), and cytolytic marker (Granzyme B), in murine CD8^+^ T cells treated with mPD-L1 EVs. Accompanying bar graphs show mean +/- s.e.m. of MFI from n=4 mice. Statistical significance was determined using ordinary one-way ANOVA, p-values are shown. Statistical significance defined as p < 0.05.

Murine CD4^+^ and CD8^+^ T cells responded to mCD80 EVs by increasing the CD25, Ki67, IL-2 and IFNγ (**Fig. 3b**). The treatment with mOX40L EVs increased CD69, Ki67, IL-2 and IFNγ in both murine CD4^+^ and CD8^+^ T cells (**Fig. 3c**) and decreased the T regulatory capacity of CD4^+^ T cells, as measured by FoxP3 (**Extended Data Fig. 3d**). As for PD-L1-containing EVs, we observed that murine CD8^+^ T cells significantly decreased their activation, cytokine production, and cytolytic potential when treated with mPD-L1 EVs (**Fig. 3d**), whereas murine CD4^+^ T cells did not undergo significant changes upon treatment with mPD-L1 EVs (**Extended Data Fig. 3c**).

Taken together, our results show that EVs harboring immune checkpoint proteins can also elicit positive and negative co-stimulation of murine T cells *ex vivo*, suggesting that this strategy can be employed to engage immune checkpoint pathways in T cells from multiple species.

### EV-driven immunomodulation alters tumor growth kinetics and remodels the immune landscape of tumors

To investigate whether the engineered EVs can modulate T cell functions in vivo and influence cancer progression, we used an orthotopic melanoma model, whereby B16-F10 cells were implanted intradermally in immunocompetent mice. The engineered EVs were administered systemically via the intraperitoneal route, and tumor growth evaluated (**Fig. 4a, e, i**). The growth rate of B16-F10 melanoma tumors was not significantly altered upon treatment with EVs harboring mCD80, although a trend towards a delay in tumor growth was observed from day 12 to 16 post-tumor implantation (**Fig. 4b-d**). Notably, the administration of EVs containing mOX40L significantly delayed the growth of B16-F10 melanoma tumors (**Fig. 4f-h**), whereas EVs harboring mPD-L1 significantly accelerated tumor growth (**Fig. 4j-l**).

**Fig. 4:**
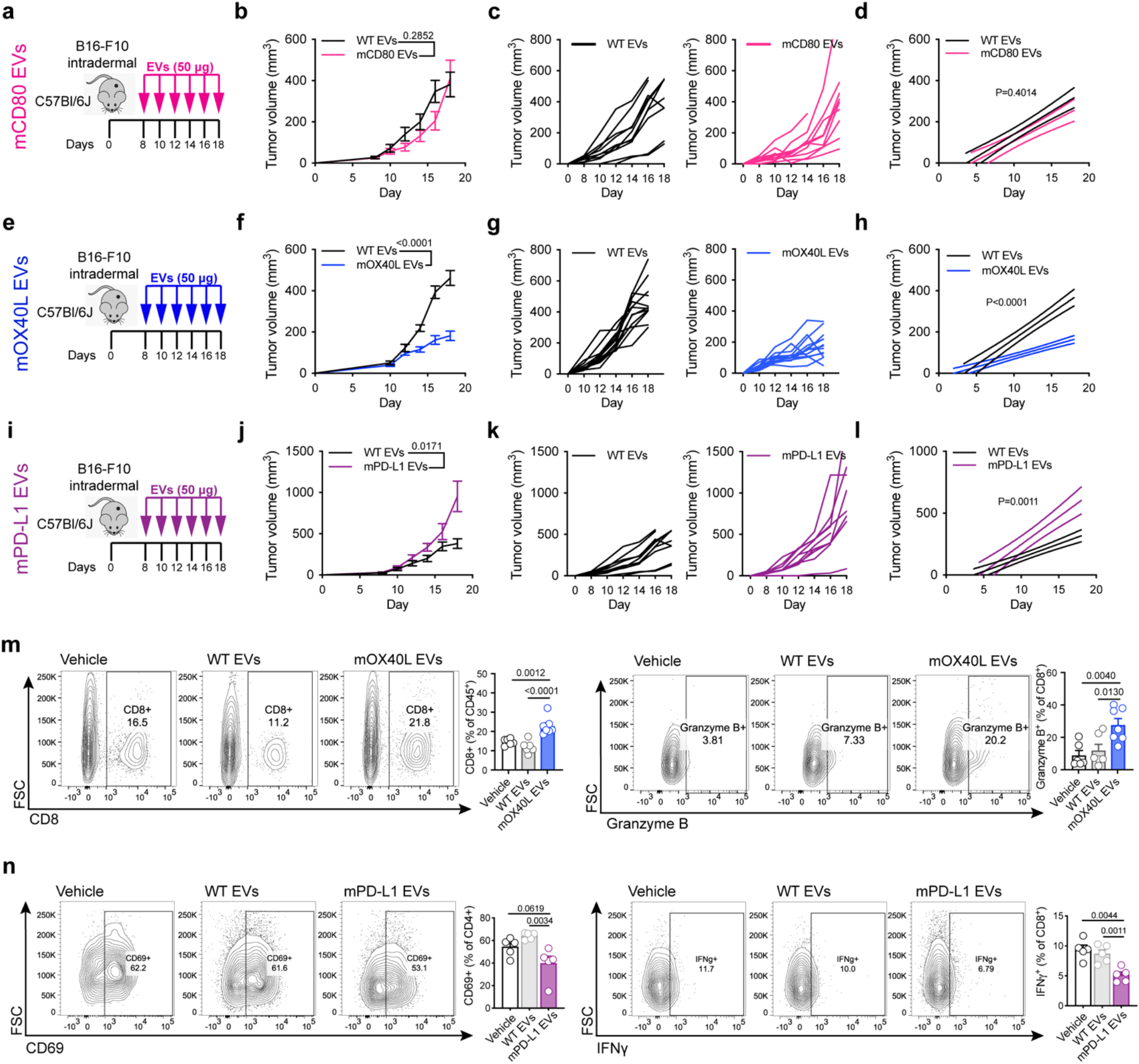
Engineered EVs modulate the growth and alter the immune landscape of melanoma tumors. (**a**) Schematic representation of experimental design employed to assess the role of mCD80 EVs in vivo. (**b**) Kinetics of B16-F10 tumor growth upon administration of WT or mCD80 EVs. Graph shows mean +/- s.e.m. of tumor volume from n=10 mice per group. Statistical significance was determined using two-way ANOVA with Šídák’s multiple comparison test. (**c**) Kinetics of B16-F10 tumor growth upon administration of WT or mCD80 EVs. Graph shows individual curves for tumor volume from n=10 mice per group. (**d**) Linear regression analysis test for significant differences in slope of tumor growth kinetics between WT and mCD80 EV treatments. Statistical significance was determined using Simple linear regression. (**e**) Schematic representation of experimental design employed to assess the role of mOX40L EVs in vivo. (**f**) Kinetics of B16-F10 tumor growth upon administration of WT or mOX40L EVs. Graph shows mean +/- s.e.m. of tumor volume from n=12-13 mice per group. Statistical significance was determined using two-way ANOVA with Šídák’s multiple comparison test. (**g**) Kinetics of B16-F10 tumor growth upon administration of WT or mOX40L EVs, graph shows individual curves for tumor volume over time from n=12-13 mice per group. (**h**) Linear regression analysis test for significant differences in slope of tumor growth curves between WT EVs and mOX40L EV treatments. Statistical significance was determined using Simple linear regression, p-value is shown. (**i**) Schematic representation of experimental design employed to assess the role of mPD-L1 EVs in vivo. (**j**) Kinetics of B16-F10 tumor growth upon administration of WT or mPD-L1 EVs. Graph shows mean +/- s.e.m. of tumor volume from n=10 mice per group. Statistical significance was determined using two-way ANOVA with Šídák’s multiple comparison test. (**k**) Kinetics of B16-F10 tumor growth upon administration of WT or mPD-L1 EVs, graph shows individual curves for tumor volume over time from n=10 mice per group. (**l**) Linear regression analysis test for significant differences in slope of tumor growth curves between WT EVs and mPD-L1 EV treatments. Statistical significance was determined using Simple linear regression, p-value is shown. Same control WT EVs used in comparison with mCD80 EVs and with mPD-L1 EVs. (**m**) The administration of mOX40L EVs promote the infiltration and the cytotoxic potential of CD8^+^ T cells. Representative contour plots show CD8^+^ T cells (left) and Granzyme B (right). Bar graphs show quantification results expressed as mean +/- s.e.m. of percentage of positive cells, n=6-7 mice per group. Statistical significance was determined using ordinary one-way ANOVA with Dunnet’s multiple comparison test. (**n**) The administration of mPD-L1 EVs reduce the activation of CD4^+^ T cells and IFNγ in CD8^+^ T cells. Representative contour plots show CD69^+^ CD4^+^ T cells (left) and IFNγ+ CD8^+^ T cells (right). Bar graphs show quantification results expressed as mean +/- s.e.m. of percentage of positive cells, n=5 mice per group. Statistical significance was determined using ordinary one-way ANOVA with Dunnet’s multiple comparison test. Statistical significance defined as p < 0.05.

To gain insights into the mechanism of action of mOX40L and mPD-L1 engineered EVs *in vivo*, we characterized the landscape of lymphocytes using multiparametric flow cytometry. We found that the systemic administration of OX40L-containing EVs altered the landscape of tumor-infiltrating lymphocytes, as measured by a significant increase in the proportion of CD3^+^CD8^+^ T cells in tumors, as well as heightened cytotoxic potential of CD8^+^ T cells, as measured by increased Granzyme B (**Fig. 4m**). In addition, the treatment with mOX40L EVs promoted increased activation, IL-2, Perforin and T-bet levels in tumor-resident CD8^+^ T cells and enhanced the proliferation of CD4^+^ T cells and CD19^+^ B cells in circulation (**Extended Data Fig. 4b**). Taken together, our findings suggest that engineered EVs harboring OX40L promote anti-tumor immunity in the B16-F10 model through enhancement of cytotoxic T lymphocyte functions. On the other hand, we observed that PD-L1-containing EVs dampened anti-tumor immunity by suppressing the activation of CD4^+^ T cells, and the production IFNγ in tumor resident CD8^+^ T cells (**Fig. 4n**). In addition, the administration of mPD-L1 EVs decreased the abundance of CD4^+^ T cells in circulation (**Extended Data Fig. 4c**).

Further, the effect of systemic administration of the engineered EVs was investigated in a second pre-clinical model, in which colorectal cancer MC-38 cells were implanted subcutaneously. In this setting, similar phenotypic changes in tumor growth kinetics were observed to the melanoma model, whereby mCD80 EVs did not significantly alter the rate of MC-38 growth, mOX40L EVs delayed MC-38 tumor growth and mPD-L1 EVs accelerated MC-38 tumor growth (**Fig. 5a-l**).

**Fig. 5:**
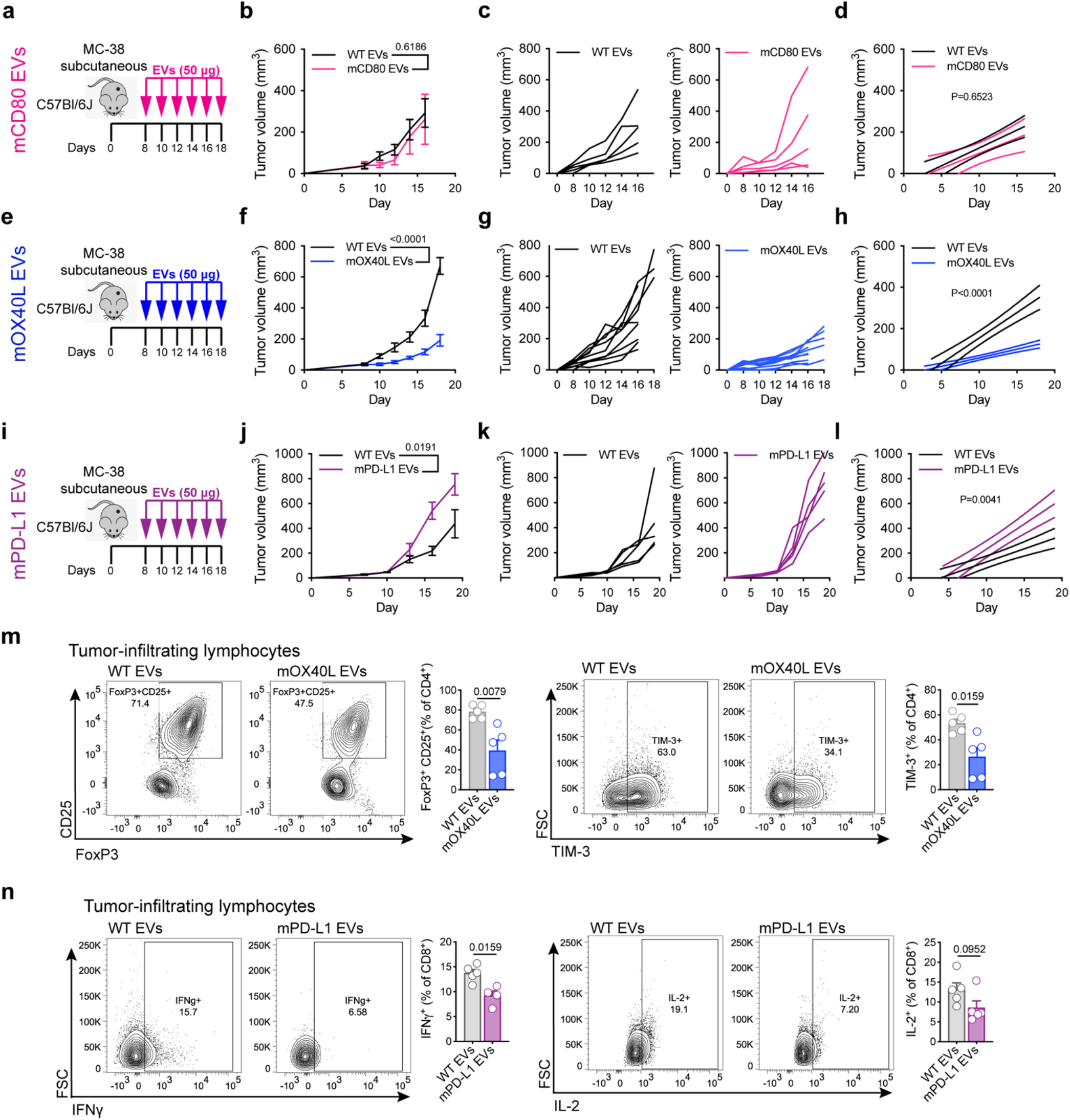
Engineered EVs modulate the growth and alter the immune landscape of colorectal tumors. (**a**) Schematic representation of experimental design employed to assess the role of mCD80 EVs in MC-38 model. (**b**) Kinetics of MC-38 tumor growth upon administration of WT or mCD80 EVs. Graph shows mean +/- s.e.m. of tumor volume from n=5 mice per group. Statistical significance was determined using two-way ANOVA with Šídák’s multiple comparison test. (**c**) Kinetics of MC-38 tumor growth upon administration of WT or mCD80 EVs. Graph shows individual curves for tumor volume from n=5 mice per group. (**d**) Linear regression analysis test for significant differences in slope of tumor growth kinetics between WT and mCD80 EV treatments in MC-38 model. Statistical significance was determined using Simple linear regression. (**e**) Schematic representation of experimental design employed to assess the role of mOX40L EVs in MC-38 model. (**f**) Kinetics of MC-38 tumor growth upon administration of WT or mOX40L EVs. Graph shows mean +/- s.e.m. of tumor volume from n=10 mice per group. Statistical significance was determined using two-way ANOVA with Šídák’s multiple comparison test. (**g**) Kinetics of MC-38 tumor growth upon administration of WT or mOX40L EVs, graph shows individual curves for tumor volume over time from n=10 mice per group. (**h**) Linear regression analysis test for significant differences in slope of tumor growth curves between WT and mOX40L EV treatments in MC-38. Statistical significance was determined using Simple linear regression. (**i**) Schematic representation of experimental design employed to assess the role of mPD-L1 EVs in MC-38. (**j**) Kinetics of MC-38 tumor growth upon administration of WT or mPD-L1 EVs. Graph shows mean +/- s.e.m. of tumor volume from n=5 mice per group. Statistical significance was determined using two-way ANOVA with Šídák’s multiple comparison test. (**k**) Kinetics of MC-38 tumor growth upon administration of WT or mPD-L1 EVs, graph shows individual curves for tumor volume over time from n=5 mice per group. (**l**) Linear regression analysis test for significant differences in slope of tumor growth curves between WT and mPD-L1 EV treatments in MC-38. Statistical significance was determined using Simple linear regression. (**m**) The administration of mOX40L EVs reduces T_regs_ and diminishes the exhaustion of CD4^+^ T cells in MC-38 tumors. Representative contour plots show FoxP3^+^ CD25^+^ T_regs_ (left) and TIM-3^+^ CD4^+^ T cells (right). Bar graphs show quantification results expressed as mean +/- s.e.m. of percentage of positive cells, n=5 mice per group. Statistical significance was determined by two-tailed unpaired Mann-Whitney t-test. (**n**) The administration of mPD-L1 EVs reduces IFNγ and IL-2 in CD8^+^ T cells. Representative contour plots show IFNγ^+^ CD8^+^ T cells (left) and IL-2^+^ CD8^+^ T cells (right). Bar graphs show quantification results expressed as mean +/- s.e.m. of percentage of positive cells, n=5 mice per group. Statistical significance was determined by two-tailed unpaired Mann-Whitney t-test. Statistical significance defined as p < 0.05.

Immunophenotypic characterization of MC-38 lymphocytes was performed using multiparametric flow cytometry. Mechanistically, the treatment of MC-38-bearing mice with mOX40L EVs decreased the proportion tumor-resident regulatory T cells (T_regs_), and reduced the exhaustion of CD4^+^ T cells, as measured by decreased TIM-3 (**Fig. 5m**). In addition, the proportion of activated versus naïve CD4^+^ T cells was altered upon administration of mOX40L EVs, whereby tumor-resident CD4^+^ T cells displayed a significantly more activated phenotype, as opposed to lymph node-resident CD4^+^ T cells, which show higher abundance of naïve CD4^+^ T cells, as measured by CD62L (**Extended Data Fig. 5a, b**). Similar to the T cell phenotypes observed in melanoma, mPD-L1 EVs reduced the production IFNγ, as well as the production of IL-2 in tumor in CD8^+^ T cells (**Fig. 5n**).

Together, using two distinct pre-clinical tumor models, our results show that EVs containing immune checkpoint proteins can elicit positive and negative co-stimulation of murine T cells, remodel the immune landscape of tumors and influence the kinetics of tumor growth.

### EV-driven immunomodulation influences the progression of acute autoimmune hepatitis

To investigate the role of immunomodulatory molecules in EVs in the context of auto-immune disorders, we exploited the Concanavalin A (ConA)-induced liver injury model, in which T cell hyperactivation causes liver damage ^30^. Immunocompetent mice were pre-treated with EVs or vehicle at 96 hours, 48 hours and immediately prior to ConA administration, tissues were harvested for analysis at 8 hours post ConA treatment (**Fig. 6a**). Histological analyses revealed that the treatment with mOX40L EVs exacerbated hepatocyte necrosis, resulting in an increased liver injury score (**Fig. 6b, c**). Mice treated with mPD-L1 EVs, and to a lesser extent, mCD80 EVs, displayed a trend towards decreased liver necrosis in comparison to control groups (**Fig. 6b, c**).

**Fig. 6:**
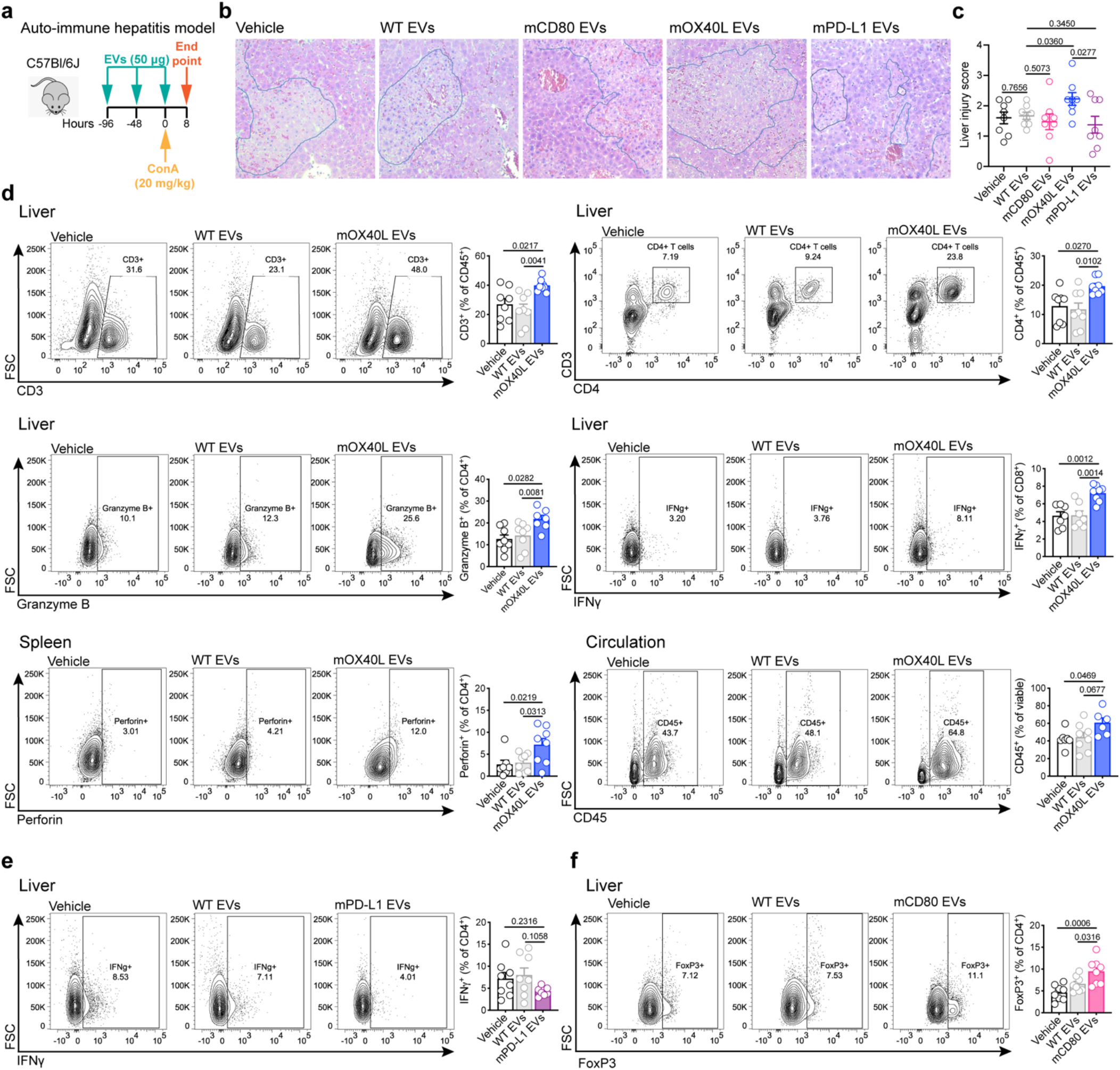
Engineered EVs modulate the severity and the immune landscape in the Concanavalin A-induced auto-immune hepatitis model. (**a**) Schematic representation of experimental design employed to assess the role of engineered EVs in the ConA-induced hepatitis model. (**b**) Images of liver sections from distinct treatment groups stained for Hematoxylin and Eosin. Highlighted areas in blue show necrotic tissue. (**c**) Histological liver injury score. Statistical significance determined by two-tailed unpaired t-test. (**d**) Immunophenotyping of liver, spleen and circulating immune cells from ConA model upon treatment with mOX40L EVs. Representative contour plots display differences in frequency of specified immune populations. Bar graphs show quantification results expressed as mean +/- s.e.m. of percentage of positive cells. Statistical significance was determined using ordinary one-way ANOVA with Dunnet’s multiple comparison test. (**e**) Immunophenotyping of liver from ConA model upon treatment with mPD-L1 EVs. Representative contour plot displays frequency of IFNγ^+^ CD4^+^ T cells. Bar graph shows quantification results expressed as mean +/- s.e.m. of percentage of positive cells. Statistical significance was determined using ordinary one-way ANOVA with Dunnet’s multiple comparison test. (**f**) Immunophenotyping of liver from ConA model upon treatment with mCD80 EVs. Representative contour plot displays frequency of FoxP3^+^ CD4^+^ T cells. Bar graph shows quantification results expressed as mean +/- s.e.m. of percentage of positive cells. Results from n=8 mice per group. Statistical significance was determined using ordinary one-way ANOVA with Dunnet’s multiple comparison test. Statistical significance defined as p < 0.05.

To further dissect the mechanism through which our engineered EVs affect the progression of autoimmune hepatitis, we profiled immune landscape of liver, spleen and blood using flow cytometry. In line with the observations of heightened histological liver injury, we observed that the liver of mice treated with mOX40L EVs displayed an increased proportion of CD3^+^ CD4^+^ T cells, with increased Granzyme B (**Fig. 6d**). In addition, we observed a higher proportion of IFNγ in CD8^+^ T cells in mice treated with mOX40L than control EVs or vehicle (**Fig. 6d**). In addition, in spleen-resident immune cells, there was an increase in Perforin^+^ CD4^+^ T cells, and in circulation, and overall increase in CD45^+^ immune cells upon treatment with mOX40L EVs (**Fig. 6d**).

The treatment with mPD-L1-containing EVs showed a trend towards decreased IFNγ in CD4^+^ T cells, while mCD80-containing EVs increased the frequency of regulatory T cells in the liver, both changes argue towards an immunosuppressive phenotype upon administration of these EVs in the ConA model (**Fig. 6e, f**).

Together, our results suggest that immunomodulatory molecules residing in EVs may condition T cells to promote or inhibit the progression of ConA-induced auto-immune hepatitis.

### Engineered EVs containing OX40L synergize with immune checkpoint blockade therapy to promote anti-tumor immunity

Based on our observations, mOX40L-containing EVs displayed an enhanced ability to delay the growth of tumors and to accelerate the progression of auto-immune hepatitis. These findings prompted us to further establish its potential as an anti-tumor therapy.

To further understand the basis of action of mOX40L EVs, we exploited the B16-F10 melanoma model. To determine the contribution of the CD4^+^ and CD8^+^ T cell subsets in the antitumor phenotype mediated by OX40L EVs in the orthotopic B16-F10 model, we depleted CD4^+^ and CD8^+^ T cells using antibodies. We validated the depletion efficacy of CD4^+^ and CD8^+^ T cell subsets using flow cytometry (**Extended Data Fig. 6**). The depletion of CD4^+^ T cells did not impair the anti-tumor efficacy of mOX40L EVs; in fact, all treatment groups receiving the anti-CD4 antibodies exhibited delayed tumor growth and extended survival, which is likely a result of depletion of regulatory T cells (T_reg_) (**Extended Data Fig. 7a-c**). On the other hand, the depletion of CD8^+^ T cells abrogated the anti-tumor effect of mOX40L EVs, as measured by the rescue in tumor growth kinetics and survival (**Extended Data Fig. 7d-f**). Together with the immunophenotyping results in the B16-F10 model, these results suggest that mOX40L EVs primary signal to CD8^+^ T cells to promote anti-tumor immunity in B16-F10 melanoma.

Clinical trials conducted in melanoma patients have shown encouraging response to immune checkpoint blockade therapy, with significant improvement in overall survival and progression free survival rates ^31^. Given the promise of harnessing the immune system to treat cancer, we sought to investigate the anti-tumor potential of mOX40L EVs as a positive co-stimulatory signal in combination with αCTLA-4 administration in orthotopically implanted B16-F10 tumors (**Fig. 7a**). Notably, we observed that the EV-mediated OX40 engagement combined with CTLA-4 blockade further delayed the growth of B16-F10 melanoma tumors and extended the survival of tumor-bearing mice in comparison to the monotherapies alone and to control groups (**Fig. 7, b-d**). Thus, our findings suggest that OX40L EVs could potentiate CTLA-4 blockade to promote antitumor immunity *in vivo* and may represent an additional avenue to treat melanoma patients.

**Fig. 7.**
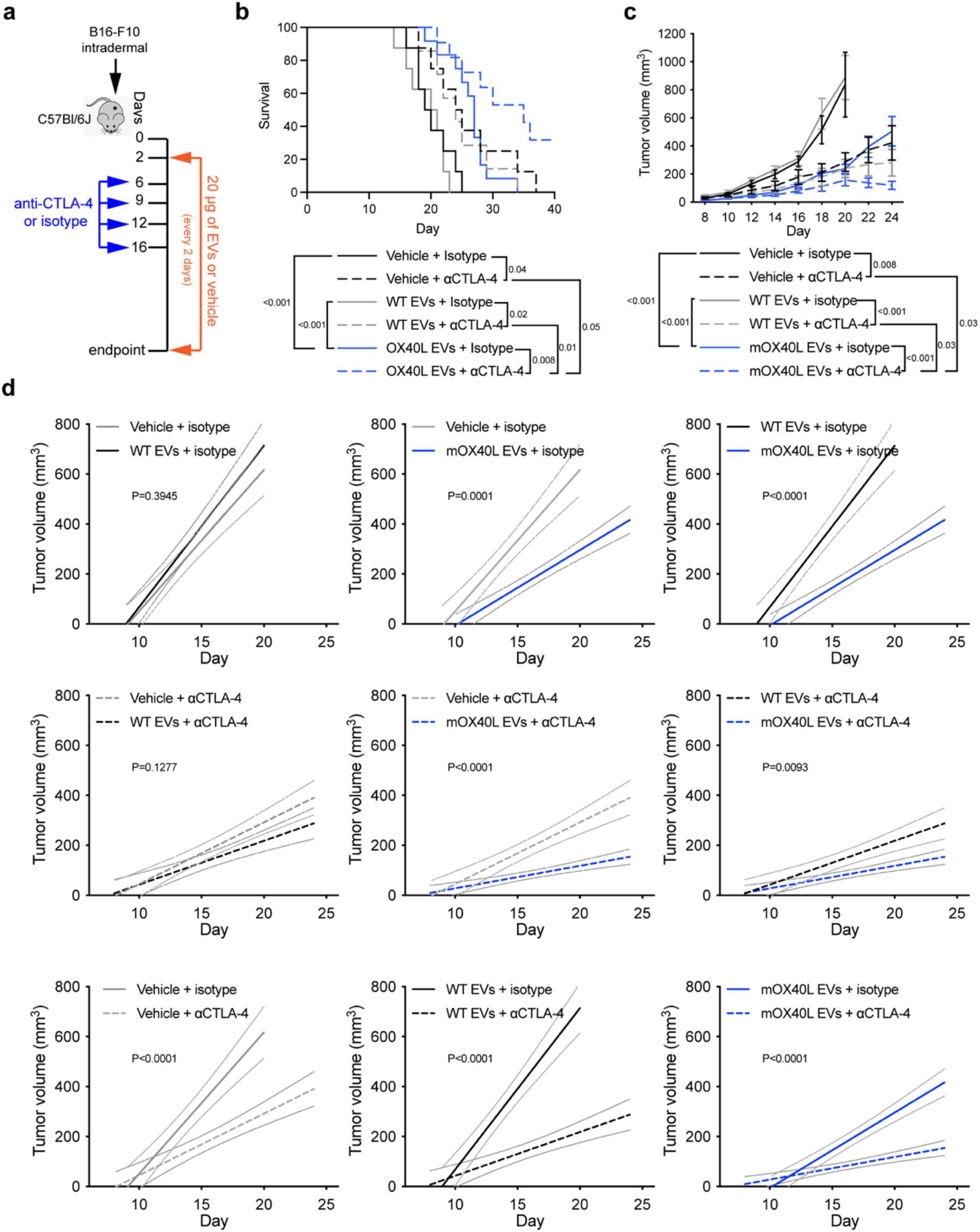
OX40L EVs synergize with anti-CTLA-4 to promote anti-tumor immunity. (**a**) Schematic representation of experimental design employed to assess the therapeutic potential of OX40L EVs combined with anti-CTLA-4 in B16-F10 melanoma. (**b**) Kaplan–Meier curve of tumor-bearing mice treated with vehicle, WT or mOX40L EVs in the presence of isotype or anti-CTLA-4 antibody. Survival plot shows results from n=8-12 mice per group. Statistical analysis performed using log-rank Mantel–Cox test. (**c**) Kinetics of B16-F10 tumor growth treated with vehicle, WT or mOX40L EVs in the presence of isotype or anti-CTLA-4 antibody. Graph shows mean +/- s.e.m. of tumor volume from n=8-12 mice per group. Statistical significance was determined using two-way ANOVA with Šídák’s multiple comparison test. (**d**) Linear regression analysis to test for significant differences in slope of tumor growth curves between experimental groups. Statistical significance was determined using Simple linear regression and p-value is shown. Statistical significance defined as p < 0.05.

## Discussion

Here, we show that epithelial EVs harboring immunomodulatory molecules can functionally elicit positive and negative T cell co-stimulation in the setting of cancer and auto-immune disease, while control EVs from epithelial cells did not elicit T cell-dependent immune responses. This suggests that native EVs can be engineered to induce adaptive immune response, showcasing their ability to engage the immune system when proper context is provided.

In the realm of cancer therapeutics, EVs have been engineered to harbor proteins that can induce apoptosis (e.g. SIRPα and TRAIL) and modulate specific cell populations of the immune system (e.g. CD40L, alarmin, CAR, IL-12 and IFNγ), approaches which have been demonstrated to pose anti-tumor properties ^18,20–23,32,33^. Our work not only adds to the existing toolkit of EV-based anti-cancer therapeutics ^17^, but also provides proof-of-principle evidence that immunomodulatory molecules in EVs can be used to alter the progression of auto-immune pathologies.

We show that EVs harboring CD80, OX40L and PD-L1 can functionally elicit positive and negative T cell co-stimulation in both human and murine CD4^+^ and CD8^+^ T cells, using *ex vivo* assays, thus suggesting applicability of these engineered EVs in the context of human pathologies. Despite displaying functional activity *ex vivo*, *in vivo* CD80 EVs did not trigger significant T cell-mediated responses in cancer and auto-immune hepatitis. This is likely because in the complex *in vivo* scenario, CD80 EVs would have the ability to bind to both CD28 and to CTLA-4, unlike to the *ex vivo* assays, in which stimulated T cells displayed preferential expression of CD28 receptor at their surface. Similar to literature reports, PD-L1 EVs efficiently accelerated the growth of B16-F10 and MC-38 tumors ^9,34^. Using flow cytometry, we found that PD-L1 EVs decreased IFNγ production by T cells. Notably, *in vivo* OX40L EVs significantly delayed the growth of B16-F10 and MC-38 tumors. In the melanoma model, we found that the mechanism of action of OX40L EVs likely occurs through CD8^+^ T cells, as demonstrated by antibody depletion and by flow cytometry, whereby OX40L EVs enhance the proportion and the cytotoxicity of CD8^+^ T cells. On the other hand, in the MC-38 model, OX40L EVs likely mediate their anti-tumor role by decreasing the proportion of regulatory T cells and the exhaustion of CD4^+^ T cells. Thus, the molecular mechanism through which OX40L EVs mediates anti-tumor responses seems to be tumor type specific. In fact, in the literature, the engagement of the OX40 pathway was demonstrated to elicit T cell responses in both CD4^+^ and CD8^+^ subsets in a tumor model-dependent manner ^35^. Advanced head and neck squamous cell carcinoma (HNSCC) patients treated with an OX40 agonist in the neoadjuvant setting, had an increased activation and proliferation of CD4^+^ and CD8^+^ T cells in circulation and tumor ^36^. Using antigen-specific OX40-deficient CD8^+^ T cells, the OX40 pathway was shown to mediate the expansion of CD8^+^ T cells and to elicit cytotoxic T lymphocyte-mediated protection against tumor growth ^37^. The engagement of OX40 with Fc-mOX40L also promoted anti-tumor immunity through CD8^+^ T cells in colon- and renal cell carcinoma-bearing mice ^38^. In addition, OX40 agonist antibody enhanced CD8^+^ T cell infiltration and reduced immunosuppression in sarcoma, mammary carcinoma, and colorectal carcinoma models ^39^. OX40-mediated co-stimulation was also shown to decrease Treg-mediated immunosuppression and to promote tumor rejection ^40,41^.

Immune checkpoint blockade therapies based on αCTLA-4 and αPD-1/αPD-L1 have revolutionized cancer treatment ^42–45^. However, clinical data shows that a long-lasting response to these treatments as single agents or as combination therapy in melanoma only occurs in a subset of cancer patients ^31^. Therefore, there is an unmet clinical need to further harness the immune system to prolong patient survival ^45^. In line with this goal, further to administering these agents as monotherapies or combination, the addition of other anti-cancer drugs has gained significant attention and several clinical trials are underway to establish their clinical value ^46^. In one of these modalities, therapeutic strategies that simultaneously combine the engagement of co-stimulatory signals in T cells with blockade of co-inhibitory signals are gaining traction ^47^. Here, we demonstrated that OX40L EVs synergize with αCTLA-4 to further delay the growth and to extend the survival of melanoma-bearing mice. Experimentally, we did not pursue the combination OX40L EVs with PD-1/PD-L1 inhibition because literature reports have shown mixed outcomes, including accelerated tumor progression ^48–50^. Of note, in human and murine melanoma tumors, αCTLA-4 and αPD-1 utilize distinct mechanisms to promote anti-tumor immunity, whereby CTLA-4 blockade induces Th1-like CD4 effector functions ^51^. Therefore, it is likely that in our study in the B16-F10 model, the synergistic anti-tumor effect observed upon OX40 engagement with CTLA-4 blockade, may occur through enhancement of cytotoxic functions mediated by OX40L EVs, with increased CD4 effector functions, promoted by αCTLA-4.

Taken together, we provide proof-of-principle evidence that immunomodulatory proteins in EVs can functionally engage T cell co-stimulation and alter the progression of cancer and auto-immune pathologies. Our study paves the way to understand the basic biology of EV-mediated positive and negative T cell co-stimulation and for further developing EV-based therapeutics that can be used in clinic.

## Materials and Methods

### Cell culture and generation of overexpressing cells

HEK293T/17 cells from ATCC (a gift from Dr. Lynda Chin’s lab), B16-F10 cells from the MD Anderson Cytogenetics and Cell Authentication Core, and MC-38 cells from Kerafast were cultured in DMEM (Corning) supplemented with 10% fetal bovine serum (FBS, Gemini) and 1% penicillin-streptomycin. Cells have been validated by STR. To generate overexpressing cells, plasmids for hCD80 (Sino Biological HG10698-UT)), hOX40L (Sino Biological HG13127-UT), hPD-L1 (Origene RC213071L4), mCD80 (Sino Biological MG50446-UT), mOX40L (Sino Biological MG53582-UT), and mPD-L1 (Origene MR203953L4) were transfected using lipofectamine 3000 (Invitrogen) in HEK293T/17 cells according to manufacturer’s instructions and selected using hygromycin (CD80 and OX40L plasmids) or puromycin (PD-L1 plasmids). Cells were tested and confirmed negative for mycoplasma and maintained in humidified cell culture incubators at 37°C and 5% CO_2_.

### EV production

Cultured cells at a confluency of approximately 80%, were washed twice with PBS and subsequently allowed to secrete EVs in serum-free medium for 24 hours (proteomics) or for 48 hours (all other assays). The conditioned medium of cells in serum-free medium was collected and subjected to differential centrifugation at 800 x g for 5 minutes and 2,000 x g for 10 minutes, and then filtered using a 0.2 μm filter flask (Corning). EVs were isolated by ultracentrifugation at 100,000 x g for 3 hours at 4°C. After ultracentrifugation, the supernatant was discarded and the EV pellet was resuspended in PBS and stored at −80°C for downstream analyses, functional assays or for in vivo treatment.

### Nanoparticle tracking analysis

The concentration and the size distribution of EVs was determined based on their Brownian motion using a NanoSight LM10 (Malvern), which is equipped with a Blue 488 nm laser and a high sensitivity sCMOS camera. The temperature was set and kept constant at 25°C during measurements. The syringe pump speed was set to 20. For each acquisition, a delay of 90 seconds followed by three captures of 30 seconds each was employed.

### RT-qPCR

RNA was isolated using RNeasy Kit (Qiagen), following manufacturer’s instructions, including the DNAse (Qiagen) treatment performed on column. RNA was resuspended in RNase-free water and the concentration was determined using Nanodrop. cDNA synthesis was performed using the High-Capacity cDNA Reverse Transcription Kit with RNase Inhibitor (Life technologies), following manufacturer’s instructions. For the qPCR reactions, Power SYBR green PCR master mix (Applied Biosystems) was used. The primers utilized are shown in **Table S1**. For data analysis, the fold-change in expression was determined using the ddCt method. Technical triplicates were used, and statistical analyses conducted on the dCt values. In the analysis of murine checkpoints overexpression, murine transcripts were not detected in human HEK293T WT cells; thus, we computed the CT values for murine CD80, OX40L, PD-L1 and of the housekeeping gene GAPDH.

### Western blot

Whole cell and EV lysates were loaded onto acrylamide gels for electrophoretic separation of proteins under denaturing conditions. Protein transfer was performed on methanol-activated PVDF membrane. Membrane was blocked in 5% BSA in TBST at room temperature for 1 hour. Antibodies are listed in **Table S2**. Visualization of immunolabels was performed with ECL solution (Pierce), following the manufacturer’s instructions. Chemiluminescent signals were captured using Amersham Hyperfilm (GE Healthcare).

### Transmission electron microscopy of EVs

EVs were fixed in 2.5% EM grade glutahaldehyde. Formvar-carbon coated mesh nickel grids were treated with poly-l-lysine solution for 5 minutes, excess was removed with filter paper and grids were allowed to dry. Drops of suspended EV samples were deposited onto the formvar-carbon coated mesh nickel grids and EVs were allowed to adsorb for 1 hour. Grids were rinsed with PBS drops 5 times, 3 minutes; and incubated in filtered 1% uranyl acetate in distilled water for 1 minute, excess was removed with filter paper and grids were allowed to dry. Grids were examined in a JEM 1010 transmission electron microscope (JEOL, USA, Inc., Peabody, MA) at an accelerating voltage of 80 kV. Digital images were obtained using AMT Imaging System (Advanced Microscopy Techniques Corp, Danvers, MA).

### Flow cytometry-based analysis of 293T cells

Approximately 5×10^5^ cells were washed in 100 μl of FACS buffer. Cell pellet was resuspended in 100 μl of FACS buffer containing either the conjugated antibody of interest or its recommended isotype control, and incubated for 30 minutes on ice protected from light. The antibodies used are listed in **Table S2.** Cells were washed three times in FACS buffer, centrifuged at 300 x g and supernatant discarded. Stained cells were resuspended in 200 μl of FACS buffer and fluorescent signal recorded with a BD LSRFortessa^TM^ X-20 equipment. Data was analyzed in FlowJo software, the percentage of positive cells was determined based on the isotype control gate of each sample (set to approximately 2%).

### Flow cytometry-based analysis of beads-bound EVs

In total, 5×10^9^ EVs were resuspended in 100 μl of PBS, and mixed to 10 μl of aldehyde/sulfate beads (Invitrogen). Samples were incubated at room temperature rotating for 15 minutes. Subsequently, 200 μl of PBS was added to each sample, vortex and incubated overnight at 4°C rotating. The following day, 150 μl of 1M glycine was added to each tube, vortex and incubated at room temperature for 1 hour rotating. The bead-bound EVs were pelleted at 13,523g for 1.5 minutes, the supernatant discarded and the precipitate resuspended in 100 μl of 10% BSA for blocking, vortex and incubated rotating at room temperature for 1 hour. After blocking, samples were centrifuged at 13,523g for 1.5 minutes, supernatant discarded and the precipitate was resuspended in 20 μl of 2% BSA containing unconjugated primary antibodies or the conjugated antibodies, alongside their corresponding isotype control, vortex and rotated at room temperature for 1 hour. After incubation with the unconjugated primary antibodies, beads-bound EVs were washed twice in 200 μl of 2% BSA, centrifuged at 13,523g for 1.5 minutes, and supernatant discarded. Beads-bound EVs were incubated for 1 hour at room temperature rotating in 20 μl of 2% BSA containing 1 μl of secondary antibody. After secondary antibody staining, or staining with conjugated antibodies, beads-bound EVs were washed three times in 200 μl of 2% BSA, centrifuged at 13,523g for 1.5 minutes, and supernatant discarded. The antibodies used are listed in **Table S2.**

After staining was finalized, the bead-bound EVs were resuspended in 500 μl of 2% BSA and fluorescent signal acquired in a BD LSRFortessa^TM^ X-20 equipment. Data was analyzed in FlowJo software, and the percentage of positive beads was determined based on the isotype control gate of each sample (set to approximately 2%).

### Ex vivo assays with human T cells

PBMCs were isolated from buffy coats using Ficoll PAQUE (GE Healthcare) and cryopreserved until further use. After thawing, PBMCs were cultured in RPMI 1640 (Corning) supplemented with 10% FBS, 1% penicillin-streptomycin, 50 μM β-mercapoethanol and 50 U/ml of human IL-2 (Peprotech) for 48 hours.

For functional assays, PBMCs were cultured in RPMI 1640 (Corning) supplemented with 10% FBS, 1% penicillin-streptomycin and 50 μM β-mercapoethanol at a cell density of 1 x 10^6^ cells/ml. T cells were activated with 1 μg/ml of anti-CD3e [OKT3] (BioLegend) and 1 μg/ml of anti-CD28 (eBioscience) for 16-24 h and then treated with EVs (50 μg/ml) or vehicle (PBS), in the presence of both anti-CD3e and anti-CD28 antibodies.

### Ex vivo assays with murine T cells

Murine spleens were dissociated through a 100 μm mesh strainer. Cells were resuspended in PBS and layered on top of Histopaque®-1119 (Sigma-Aldrich), centrifuged at 700 x g for 15 min at room temperature, with the brake set to 0. Cells from the intermediary phase were collected and washed three times in PBS. Splenocytes were cultured in RPMI 1640 (Corning) supplemented with 10% FBS, 1% penicillin-streptomycin and 50 μM β-mercapoethanol at a cell density of 1 x 10^6^ cells/ml. T cells were activated with 1 μg/ml of anti-CD3e (BD) and 1 μg/ml of anti-CD28 (BD) for 16-24 hours and then treated with EVs (50 μg/ml) or vehicle (PBS) in the presence of both anti-CD3e and anti-CD28 functional antibodies.

### Flow cytometry-based analysis of T cells from ex vivo assays

The staining procedure was performed protected from light. Cells were washed in FACS buffer, resuspended in surface staining mix containing FACS buffer with 1/2 of final volume of Brilliant Stain Buffer (BD), 50 μg/ml anti-mouse CD16/CD32 (2.4G2) (Tonbo Biosciences), surface antibodies and Fixable Viability Dye eFluor 780 (eBioscience), and incubated for 1 hour on ice. Subsequently, cells were washed 3 times in FACS buffer and incubated with Cytofix/Cytoperm (BD) for 1 hour on ice. Cells were washed 3 times with 1x perm/wash buffer (BD), and stained with intracellular staining mix containing intracellular antibodies in 1x perm/wash buffer (BD), and incubated for 1 hour on ice. The antibodies used are listed in **Table S2**. After staining, cells were washed 3 times with 1x perm/wash buffer (BD), and fixed using Cytofix fixation buffer (BD). Samples were run on a BD LSRFortessa^TM^ X-20 equipment and the data was analyzed using FlowJo software.

### Sample preparation for mass spectrometry

Clear lysates from cells and EVs in 8M urea buffer (8 M urea, 75 mM NaCl, 50 mM Tris) were quantified using microBCA Protein Assay Reagent Kit (Thermo Scientific) following the manufacturer’s instructions. For in-solution digestion, 10 μg of protein lysates derived from cells or EVs were used. Proteins were reduced using 1 mM DTT (Sigma) for 1 hour at room temperature, and alkylated using 5.5 mM of IAA (Sigma) for 45 minutes, at room temperature protected from light. Samples were diluted 8-fold in 50 mM ammonium bicarbonate (Sigma). To cleave the proteins into peptides, Trypsin/LysC mix mass-spec grade (Promega) was used at 1:10 enzyme:protein ratio, for an overnight digestion at room temperature. In the following day, samples were acidified to pH <4.0 using 1% TFA (Sigma). Peptides were desalted and concentrated using Pierce C18 spin tips (Thermo Scientific), following manufacturer’s instructions.

### nLC-MS/MS

Digested peptides were run on a Q-Exactive HF mass spectrometer coupled to an EASY-nLC II 1200 chromatography system (Thermo Scientific). Samples were loaded on a 50 cm fused silica emitter that was packed in-house with ReproSIL-Pur C18-AQ, 1.9 µm resin. The emitter was heated to 50°C using a column oven (Sonation). Peptides were eluted at a flow rate of 300 nl/minute for 125 minutes using a two-step gradient of solvent B 80% Acetonitrile:0.1% Formic Acid 2-20% in 73 minutes and to 41% at 93 minutes. Peptides were injected into the mass spectrometer using a nanoelectropsray ion source (Thermo Scientific) coupled with an Active Background Ion Reduction Device (ABIRD, ESI Source Solutions) to decrease air contaminants. Data was acquired with the Xcalibur software (Thermo Scientific) in positive mode using data-dependent acquisition. The full scan mass range was set to 375-1500m/z at 60,000 resolution. Injection time was set to 20 milliseconds with a target value of 3E6 ions. HCD fragmentation was triggered on the 15 most intense ions for MS/MS analysis. MS/MS injection time was set to 75 milliseconds with a target of 5E4 ions and with a resolution of 15,000. Ions that have already been selected for MS/MS were dynamically excluded for 13 seconds.

### MS data processing and analysis

The raw MS files were processed in the MaxQuant software ^52^ version 1.6.1.0. Andromeda search engine ^53^ performed the search against the human UniProt database. For identification, a false discovery rate (FDR) of 1% was used as threshold at both the peptide and the protein levels. Analysis settings included: a minimal peptide length of 7 amino acids, specificity for trypsin and maximum two missed cleavages. For label-free quantification, the LFQ parameter was enabled. For modifications: Acetyl (protein N-term) and oxidation (M) were set as variable modifications, and carbamidomethyl (C) as fixed modification. The match between runs parameter was not enabled. The remainder parameters used were MaxQuant default settings.

The output table proteinGroups.txt from MaxQuant was loaded into Perseus software ^54^ version 1.6.6.0 for downstream analyses. Data was filtered to remove potential contaminants, reverse peptides that match a decoy database, and proteins only identified by site. For unambiguous identification, proteins with at least one unique peptide were used for analyses. The LFQ intensities were log_2_-transformed. The summed intensities were log_10_-transformed. Annotations from Gene Ontology Cellular Compartment (GOCC), Gene Ontology Biological Processes (GOBP), Gene Ontology Molecular Function (GOMF) and Kyoto Encyclopedia of Genes and Genomes (KEGG) were added.

### In vivo studies

Female C57BL/6J mice were purchased from The Jackson Laboratory. Mice were housed in ventilated cages on a 12 hour light:12 hour dark cycle at 21–23°C and 40–60% humidity. Mice had access to an irradiated diet and sterilized water *ad libitum*.

For tumor models, implantation of 5×10^4^ B16-F10 cells resuspended in 100 μl of PBS was performed intradermally into the left flank of mice. Implantation of 1×10^5^ MC-38 cells resuspended in 100 μl of PBS was performed subcutaneously into the left flank of mice. For experiments using EVs administration as single treatment, injections started at day 8 post tumor implantation. Vehicle (PBS) or EVs (50 μg) in 100 μl of PBS were administered IP every other day. For the experiment combining EVs administration with CD4 depletion and CD8 depletion, vehicle (PBS) or EVs (50 μg) in 100 μl of PBS were administered IP every other day starting from day 8 post tumor implantation until endpoint. Isotype control or depletion antibodies were administered IP from day 6 to day 30 post-tumor implantation in 100 μl of PBS. The dosage from day 6 to 20 was of 200 μg/mouse, and from day 20 to 30 was of 100 μg/mouse. For the experiment combining EVs administration with anti-CTLA-4, vehicle (PBS) or EVs (20 μg of WT or 20 μg of mOX40L EVs) in 100 μl of PBS were administered IP every other day starting from day 2 post tumor implantation until endpoint. Isotype control or anti-CTLA-4 were administered IP at days 6, 9, 12, and day 16 post-tumor implantation in 100 μl of PBS. The dosage at day 6 was of 200 μg/mouse, and at days 9, 12 and 16 of 100 μg/mouse. The antibodies used are listed in **Table S2**. For all experiments, the tumor dimensions were measured using a digital caliper, and the volume was calculated using the formula V (mm^3^) = 0.52*a*b^2 (a=length and b=width).

For ConA-induced auto-immune hepatitis model, mice were pre-treated with EVs or vehicle IP at −96 hours, −48 hours and immediately prior to 20 mg/kg Concanavalin A (Sigma) intravenous administration at 0 hour. The tissues were harvested for analysis 8 hours post ConA treatment.

### Immunophenotyping of lymphocytes from murine tissues using flow cytometry

Immune cells from tumors, spleen, blood, liver and inguinal lymph nodes were isolated to characterize the distinct lymphocytes subsets by flow cytometry.

Tumors and liver were cut is small pieces, dissociated using gentleMACS (Miltenyi Biotec) and digested in a solution containing DNase I (Roche) and Liberase TL (Roche) in RPMI 1640 media for 30 minutes, 150 rpm, at 37°C. After digestion, tumors were further dissociated using gentleMACS (Miltenyi Biotec) and strained through a 100 μm mesh strainer. Cells were washed in PBS, resuspended in PBS and layered on top of Histopaque (Sigma-Aldrich), spin at 700 x g for 20 minutes at room temperature with the brake set to 0. Cells from the intermediary phase were collected and washed in PBS. Immune cells from spleen and inguinal lymph nodes were dissociated through a 100 μm mesh strainer. Circulating immune cells were isolated from peripheral blood. Red blood cell lysis was performed using ACK lysis buffer. After isolation, all cell populations were washed once in PBS and twice in FACS buffer.

The staining procedure was performed protected from light. Immune cells were resuspended in surface staining mix containing FACS buffer with 1/2 of final volume of Brilliant Stain Buffer (BD), 50 μg/ml anti-mouse CD16/CD32 (2.4G2) (Tonbo Biosciences), surface antibodies and Fixable Viability Dye eFluor 780 (eBioscience), and incubated for 1 hour on ice. Subsequently, cells were washed 3 times in FACS buffer and fixed/permeabilized using Fixation/Permeabilization reagent (eBioscience), following manufacturer’s instructions. Cells were washed 3 times with 1 x Permeabilization buffer (eBioscience) and stained with intracellular staining mix containing intracellular antibodies in 1 x Permeabilization buffer (eBioscience), incubated for 1 hour on ice. Cells were washed 3 times with 1 x Permeabilization buffer (eBioscience) and fixed using Cytofix fixation buffer (BD). The antibodies used are listed in **Table S2**. Samples were run on a BD LSRFortessa^TM^ X-20 equipment and the data analyzed using FlowJo software.

### Liver histopathological assessment

To evaluate the histology of the livers of mice with ConA induced hepatitis and treated with various immunomodulating EVs. Formalin-fixed paraffin embedded livers were sectioned at 5μm thickness and stained them with hematoxylin & eosin (H&E). H&E-stained livers were imaged at 200X and randomly selected 5 visual fields per mouse liver. Areas of necrosis, apoptosis and damaged hepatocytes in liver parenchyma were quantified according to the following grading scale: 0; no hepatocyte damage, no necrosis or apoptosis, 1; total areas of hepatocyte damage, necrosis and apoptosis occupy 1 to 25% of each visual field, 2; total areas of hepatocyte damage, necrosis and apoptosis occupy 26 to 50%, 3; total areas of hepatocyte damage, necrosis and apoptosis occupy 51 to 75%, 4; total areas of hepatocyte damage, necrosis and apoptosis occupy 75 to 99%, 5; total areas of hepatocyte damage, necrosis and apoptosis occupy 100%. An arithmetic mean of 5 visual fields for each mouse were calculated, which represents the liver injury score.

### Statistical analysis

Statistical analyses were performed using GraphPad Prism Software and the specific tests used are indicated in the figure legends. Unless otherwise stated, data is presented as mean values with standard error of mean (SEM). For the proteomics experiments, statistical analyses were conducted using Perseus software.

## Acknowledgments

We thank Dunner Jr. and the High-resolution Electron Microscopy Facility at the MD Anderson Cancer Center (CCSG grant NIH P30CA016672) for assistance with TEM experiments, the staff at the Cytogenetics and Cell Authentication Core at MD Anderson Cancer Center for STR testing. We thank Laura Snowden and Michelle Kirtley for technical and administrative support. F.G.K. was supported by a Post-Doctoral Training Fellowship from The Odyssey Program and Theodore N. Law for Scientific Achievement at The University of Texas MD Anderson Cancer Center. K.M.M. was supported by an Ergon Foundation Post-Doctoral Trainee Fellowship. This work in R.K. laboratory was supported with research funds from The University of Texas MD Anderson Cancer Center and NIH R35CA263815 and R01CA231465. This work was supported by funds from MD Anderson Cancer Center to RK and by a gift from Fifth Generation, Inc. (“Love, Tito’s”) and Black Rhino family office to the Kalluri Laboratory at the University of Texas MD Anderson Cancer Center. S.Z. funding is supported by Cancer Research UK (A29800).

## Author contributions

Project idea generation and planning: RK

Experimental conceptualization: FGK, VSL, RK.

Methodology: FGK, HS, DPD, YF, KH, SL, KMM, VSL, SZ.

Investigation: FGK, HS, DPD, KAA, YF, LH, KH, SL, KMM.

Visualization: FGK, HS.

Supervision: RK, SZ.

Writing: FGK, RK.

## Data and materials availability

The data supporting the findings of this study are available in the manuscript or as supplementary data. The raw files from MS analysis of cells and EVs, and the search/identification files obtained with MaxQuant were deposited to the ProteomeXchange Consortium^55^ through the PRIDE^56^ partner repository, accession number PXD044115. Username: Password: J9sG2feF

---

**Extended Data Fig. 1.**
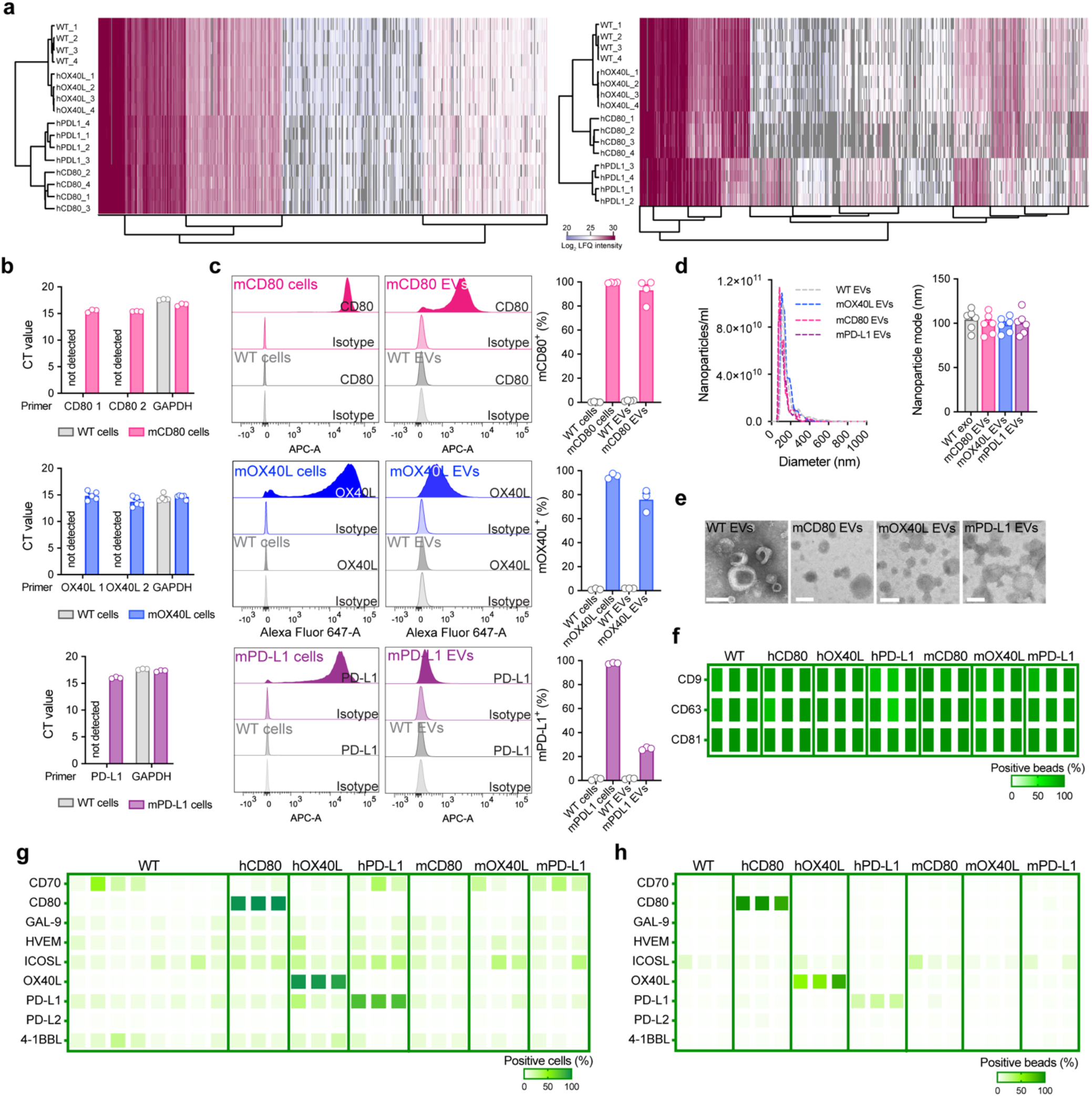
EVs can be modified to harbor high levels of the immunomodulatory checkpoint proteins CD80, OX40L and PD-L1. (**a**) Hierarchical clustering of proteomes from WT and engineered cells (left) and EVs (right) using label-free MS-based proteomics, results from n=4 biological replicates. (**b**) Expression of murine CD80, *OX40L and CD274* (PD-L1) in parental cells determined by RT-qPCR. Bar graph shows mean +/- s.e.m. of CT values from n=3-5 biological replicates. Expression of *GAPDH* used as housekeeping gene. (**c**) Flow cytometry-based evaluation of overexpressing murine proteins at the surface of parental cells and EVs. Overlaid histograms for each protein show the profile for WT and engineered cells and EVs stained with isotype control and with the antibody of interest. The accompanying bar graphs show mean +/- s.e.m. of percentage of positive cells and of positive beads (for EV analyses). Individual data points from n=3-4 biological replicates are shown. (**d**) Representative NTA profile of WT and engineered EVs with murine proteins. Bar graph shows mean +/- s.e.m. of nanoparticle mode measurements from n=6 biological replicates. (**e**) Morphology of WT and engineered EVs with murine proteins determined by TEM. Scale bar = 100 nm. (**f**) On-beads flow cytometry-based evaluation of EV surface markers CD9, CD63 and CD81 in WT and engineered EVs. Heatmap shows percentage of positive beads. Results from n=3 biological replicates. (**g**) Flow cytometry-based evaluation of co-stimulatory (human OX40L, iCOSL, 4-1BBL, CD70), co-inhibitory (human Gal-9, HVEM, PD-L1, PD-L2) and dual-role (human CD80) immune checkpoint proteins at the surface of parental cells. Results from n=3-8 biological replicates. (**h**) Flow cytometry-based evaluation of co-stimulatory (human OX40L, iCOSL, 4-1BBL, CD70), co-inhibitory (human Gal-9, HVEM, PD-L1, PD-L2) and dual-role (human CD80) immune checkpoint proteins at the surface of EVs on beads. Results from n=3 biological replicates.

**Extended Data Fig. 2.**
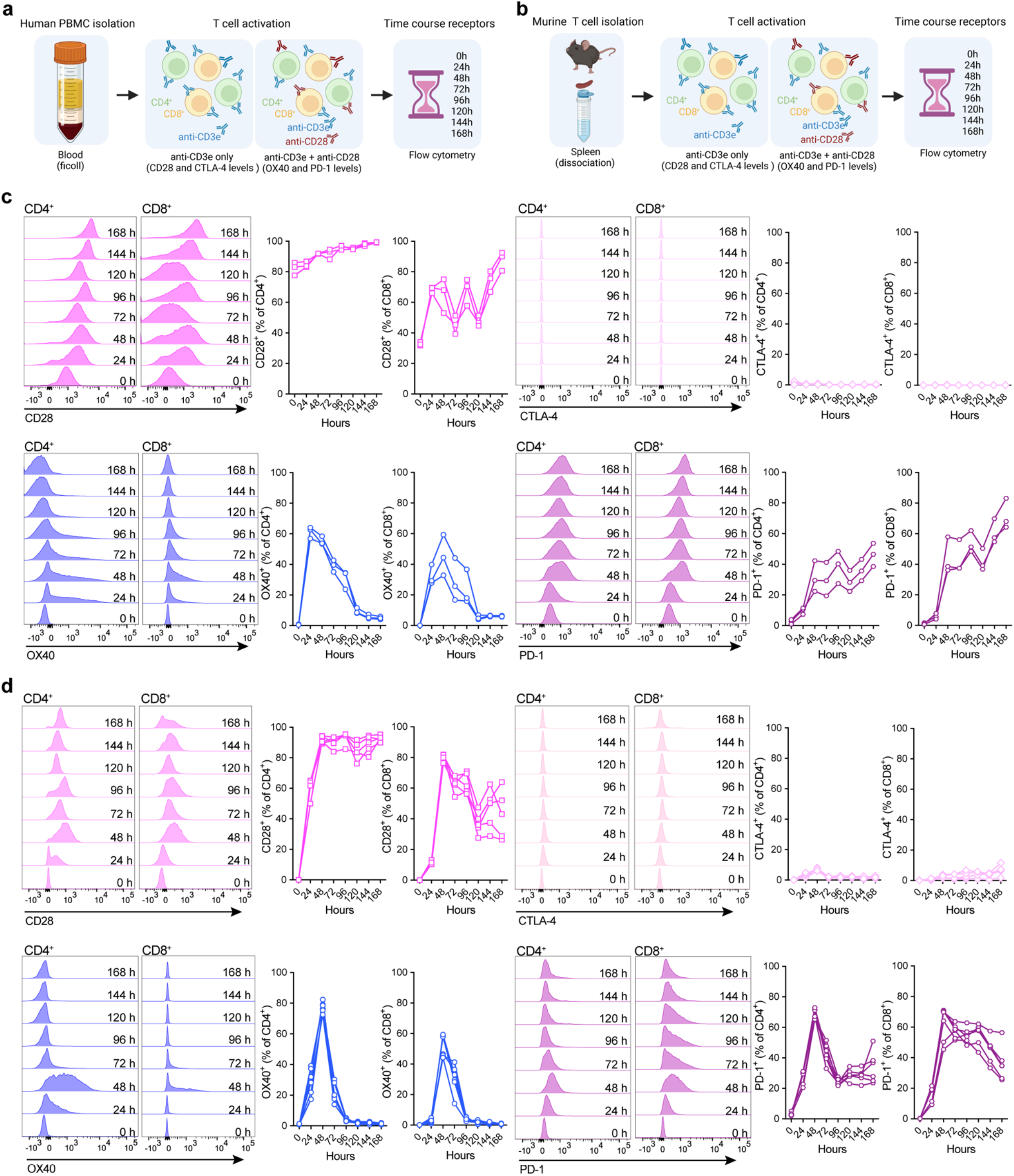
Time course evaluation of CD28, CTLA-4, OX40 and PD-1 receptors in human and murine T cells. (**a**) Schematic representation of the experimental design employed to investigate the receptors surface levels over time in the *ex vivo* assays with human PBMCs. (**b**) Schematic representation of the experimental design deployed to investigate the receptors kinetics over time in the *ex vivo* assays with murine splenocytes. (**c**) Representative overlaid histograms of human CD28, CTLA-4, OX40 and PD-1 receptors kinetics in CD4^+^ and CD8^+^ T cells. The accompanying quantification plots show the receptors levels over time in n=3 samples. (**d**) Representative overlaid histograms of murine CD28, CTLA-4, OX40 and PD-1 receptors kinetics in CD4^+^ and CD8^+^ T cells. The accompanying quantification plots show the receptors levels over time in n=5 mice.

**Extended Data Fig. 3:**
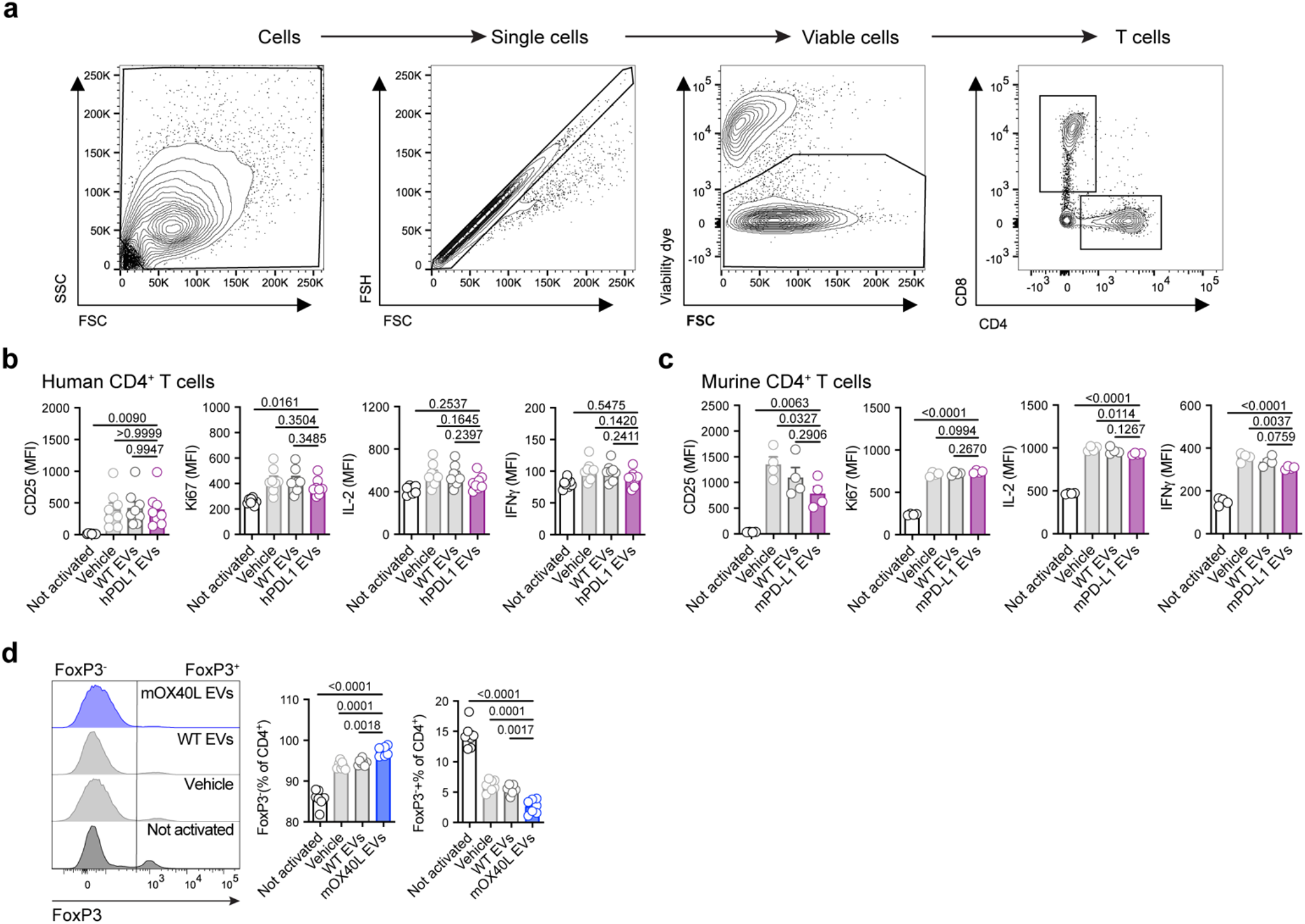
*Ex vivo* assays with engineered EVs. (**a**) Pre-gating strategy used in human and mouse *ex vivo* T cell assays. (**b**) Flow cytometry-based analysis of activation marker (CD25), proliferation marker (Ki67), and cytokines (IL-2 and IFNγ), in human CD4^+^ T cells treated with hPD-L1 EVs. Accompanying bar graphs show mean +/- s.e.m. of MFI from n=8 PBMC samples. (**c**) Flow cytometry-based analysis of activation marker (CD25), proliferation marker (Ki67), and cytokines (IL-2 and IFNγ), in murine CD4^+^ T cells treated with mPD-L1 EVs. Bar graphs show mean +/- s.e.m. of MFI from n=4 mice. (**d**) Flow cytometry-based analysis of FoxP3, in murine CD4^+^ T cells treated with mOX40L EVs. Accompanying bar graphs show mean +/- s.e.m. of MFI from n=7 mice. Statistical significance was determined using ordinary one-way ANOVA. Statistical significance defined as p < 0.05.

**Extended Data Fig. 4:**
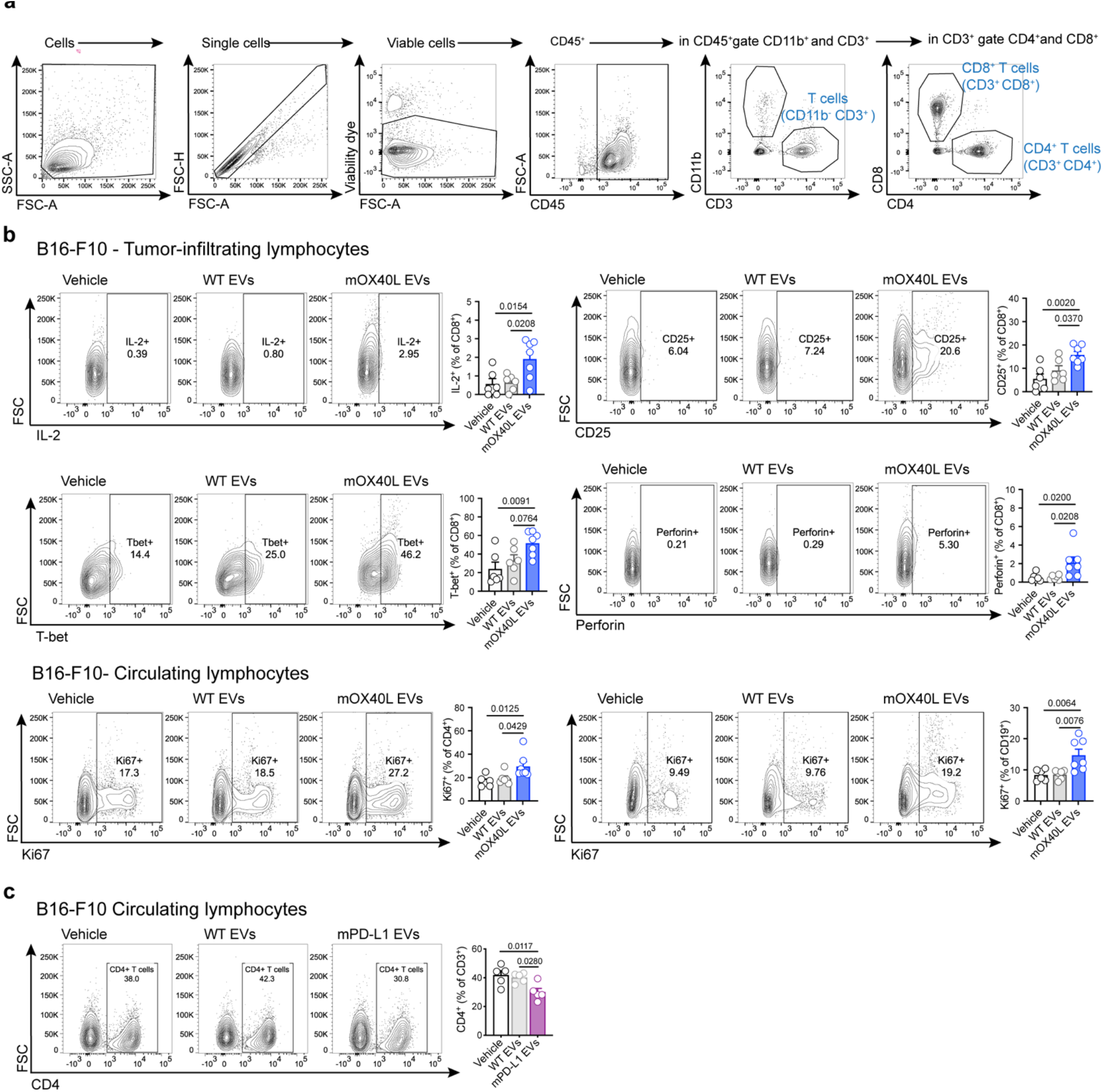
Engineered EVs alter the immune e landscape of melanoma tumors. (**a**) Pre-gating strategy used to define murine CD4^+^ and CD8^+^ T cells from *in vivo* experiments. (**b**) The administration of mOX40L EVs enhance the proportion of IL-2^+^, CD25^+^, T-bet^+^ and Perforin^+^ in tumor-infiltrating CD8^+^ T cells, and of proliferating CD4^+^ T cells and CD19^+^ B cells in circulation. Representative contour plots show the levels of each marker in the different treatment groups. Bar graphs show quantification results expressed as mean +/- s.e.m. of percentage of positive cells, n=6-7 mice per group. Statistical significance was determined using ordinary one-way ANOVA with Dunnet’s multiple comparison test. (**c**) The administration of mPD-L1 EVs reduce the proportion of CD4^+^ T cells in circulation. Representative contour plots of CD4^+^ T cells. Bar graph shows quantification results expressed as mean +/- s.e.m. of percentage of positive cells, n=5 mice per group. Statistical significance was determined using ordinary one-way ANOVA with Dunnet’s multiple comparison test. Statistical significance defined as p < 0.05.

**Extended Data Fig. 5:**
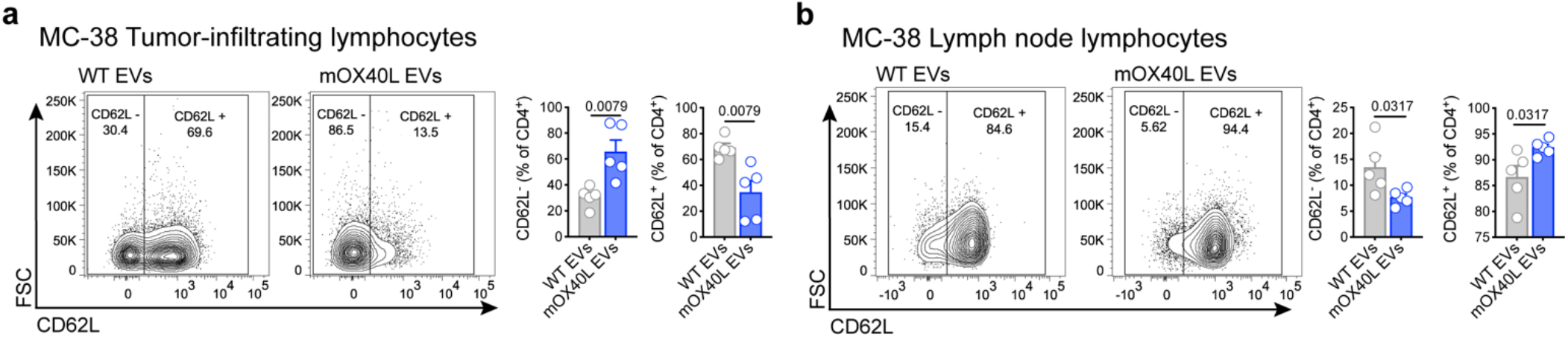
Engineered EVs alter the immune landscape of MC-38 tumor-bearing mice. The administration of mOX40L EVs alters the proportion of naïve (CD62L^+^) versus activated (CD62L^-^) in (**a**) tumor-infiltrating and in (**b**) lymph node-resident CD4^+^ T cells. Representative contour plots show the CD62L^+^ and CD62L^-^ populations in the distinct treatment groups. Bar graphs show quantification results expressed as mean +/- s.e.m. of percentage of positive and negative cells for CD62L, n=5 mice per group. Statistical significance was determined by two-tailed unpaired Mann-Whitney t-test. Statistical significance defined as p < 0.05.

**Extended Data Fig. 6.**
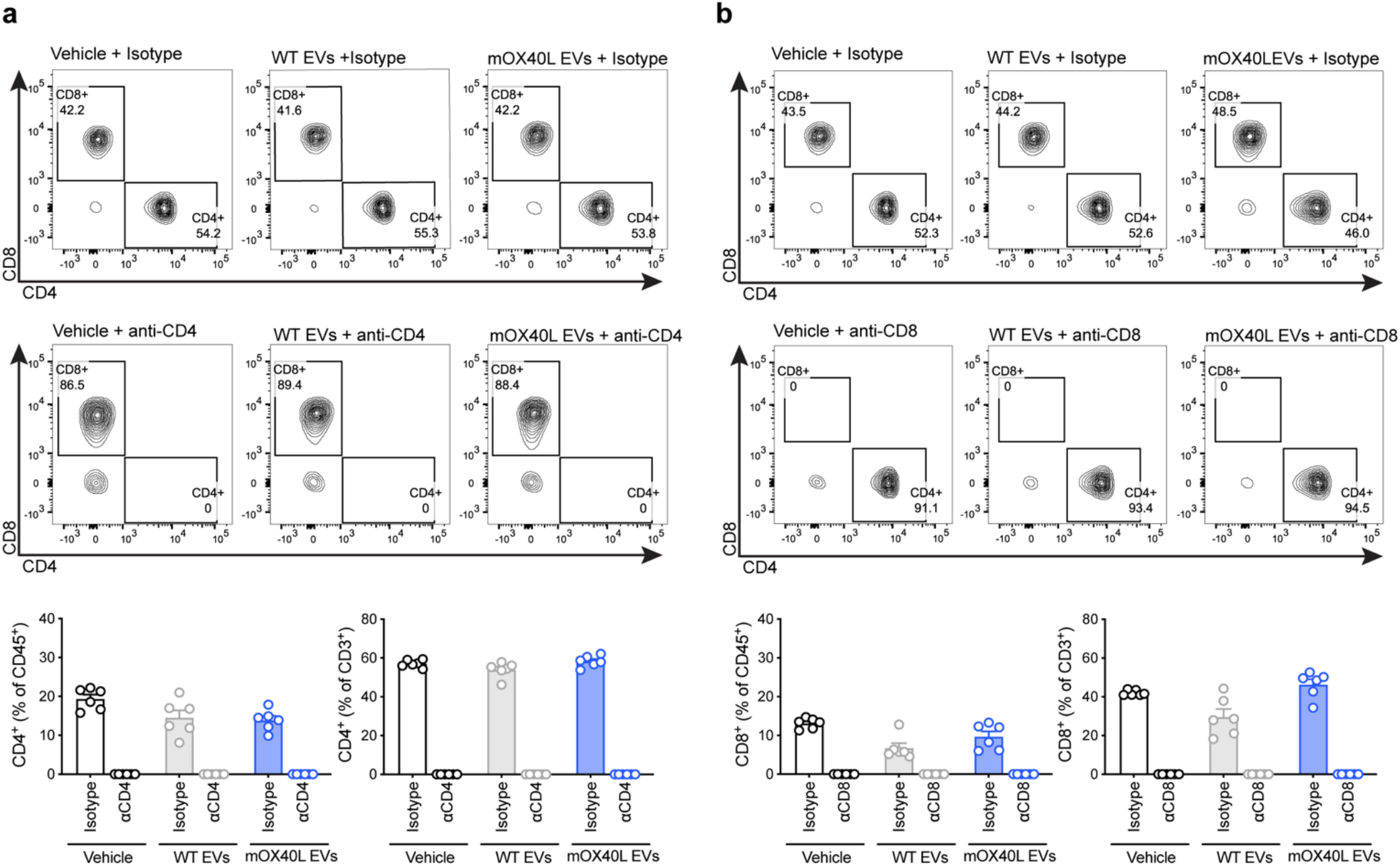
Validation of CD4 and CD8 depletion *in vivo*. (**a**) Representative contour plots of CD4^+^ and CD8^+^ T cells in the circulation of tumor-bearing mice treated with vehicle, WT or mOX40L EVs in the presence of isotype or anti-CD4 antibody. Accompanying bar graphs show quantification results expressed as mean +/- s.e.m. of percentage of CD4 positive cells, individual data points from n=6 mice per group are shown. (**b**) Representative contour plots of CD4^+^ and CD8^+^ T cells in the circulation of tumor-bearing mice treated with vehicle, WT or mOX40L EVs in the presence of isotype or anti-CD8 antibody. Accompanying bar graphs show quantification results expressed as mean +/- s.e.m. of percentage of CD8 positive cells, individual data points from n=6 mice per group are shown.

**Extended Data Fig. 7.**
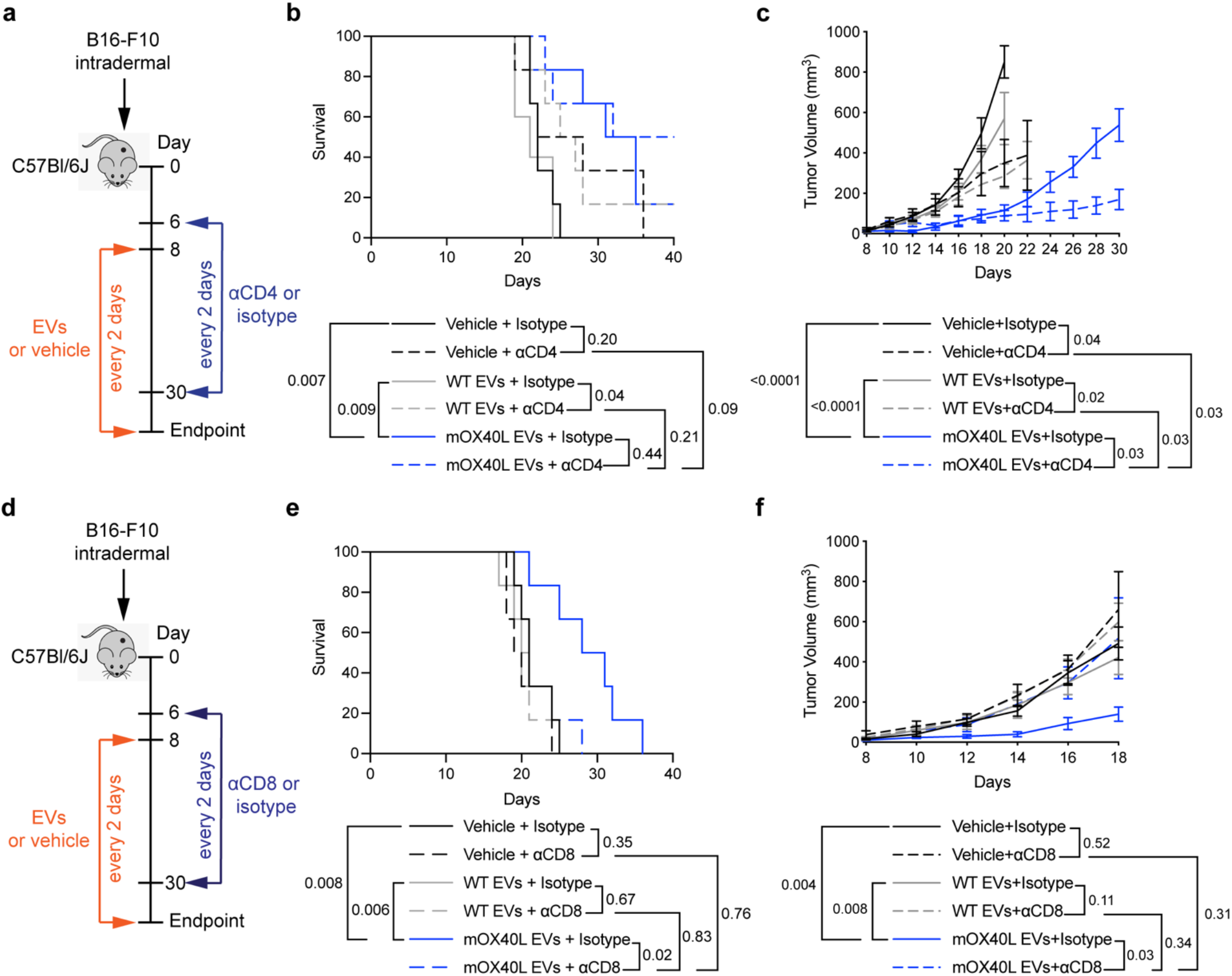
OX40L EVs promotes anti-tumor immunity in B16-F10 melanoma in a CD8^+^ T cell-dependent manner. (**a**) Schematic representation of experimental design employed to assess the contribution of CD4^+^ T cells in the anti-tumor phenotype elicited by OX40L EVs, using anti-CD4 antibody. (**b**) Kaplan–Meier curve of tumor-bearing mice treated with vehicle, WT or mOX40L EVs in the presence of isotype or anti-CD4 antibody. Survival plot shows results from n=6 mice per group. Statistical analysis performed using log-rank Mantel–Cox test. (**c**) Kinetics of B16-F10 tumor growth treated with vehicle, WT or mOX40L EVs in the presence of isotype or anti-CD4 antibody. Graph shows mean +/- s.e.m. of tumor volume from n=6 mice per group. Statistical significance was determined using two-way ANOVA with Šídák’s multiple comparison test. (**d**) Schematic representation of experimental design employed to assess the contribution of CD8^+^ T cells in the anti-tumor phenotype elicited by OX40L EVs, using anti-CD8 antibody. (**e**) Kaplan–Meier curve of tumor-bearing mice treated with vehicle, WT or mOX40L EVs in the presence of isotype or anti-CD8 antibody. Survival plot shows results from n=6 mice per group. Statistical analysis performed using log-rank Mantel–Cox test. (**f**) Kinetics of B16-F10 tumor growth upon treatment with vehicle, WT or mOX40L EVs in the presence of isotype or anti-CD8 antibody. Graph shows mean +/- s.e.m. of tumor volume from n=6 mice per group. Statistical significance was determined using two-way ANOVA with Šídák’s multiple comparison test. Statistical significance defined as p < 0.05.

**Extended Data Fig. 8.**
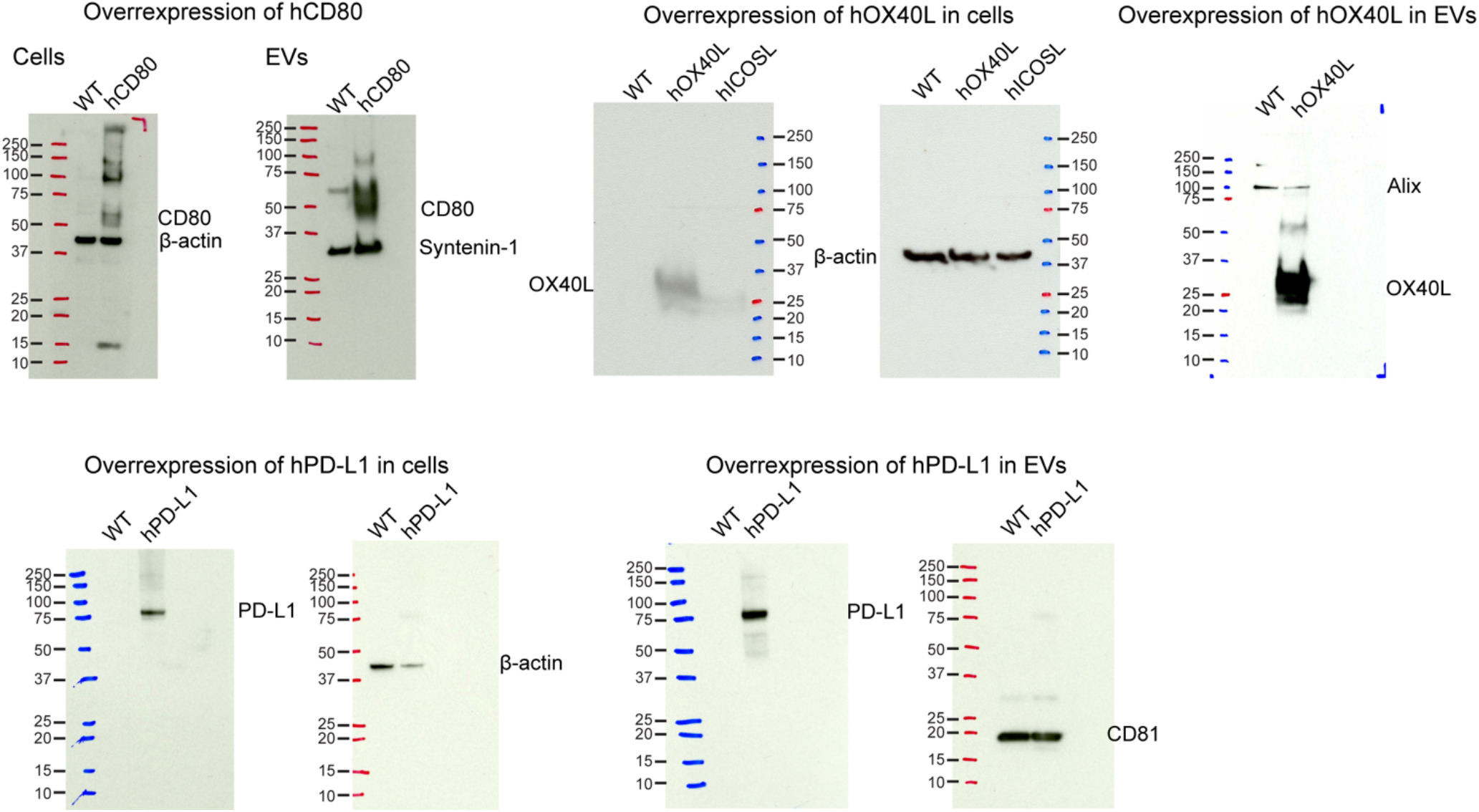
Full blots depicted in figure 1.

**Table S1:**
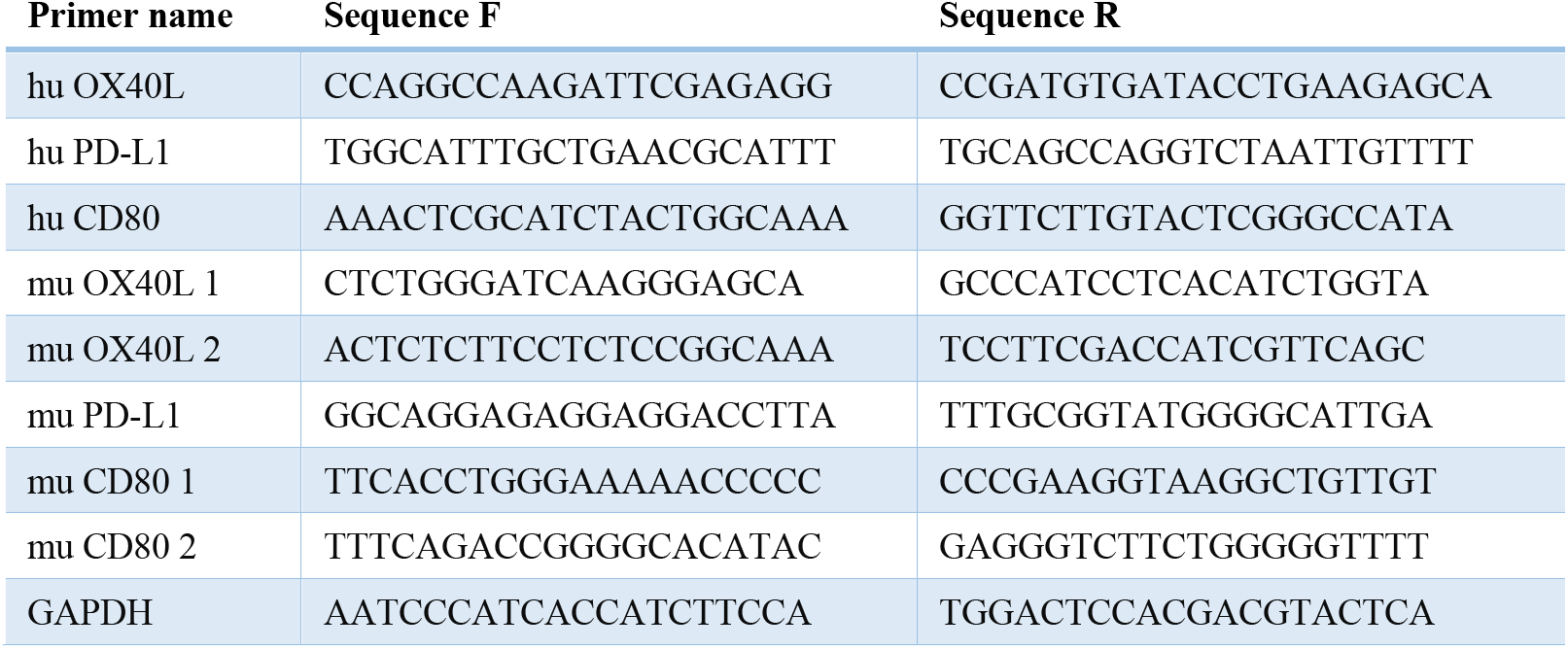
Primers.

**Table S2:** Antibodies.

| Antibody [clone] | Provider and Catalogue # | Concentration | Use |
| --- | --- | --- | --- |
| OX40L [D6X2D] | CST #14991 | 1 in 1,000 | Western blot |
| PD-L1 [MIH5] | Novus Biologicals #NBP1-43262 | 1 in 500 | Western blot |
| CD80 | Abcam # ab254579 | 1 in 500 | Western blot |
| β-Actin [13E5] | CST #4970 | 1 in 1,000 | Western blot |
| Alix [3A9] | CST #2171S | 1 in 1,000 | Western blot |
| CD81 [B-11] | Santa Cruz #sc166029 | 1 in 1,000 | Western blot |
| Syntenin-1 [EPR8102] | Abcam #ab133267 | 1 in 2,000 | Western blot |
| anti-rabbit HRP-conjugated | CST #7074 | 1 in 5,000 | Western blot |
| anti-mouse HRP-conjugated | R&D #HAF007 | 1 in 1,000 | Western blot |
| IgG1κ isotype [MOPC-21] | BD #555746 | 0.2 µg per staining | Flow cytometry (beads assay) |
| CD81 [JS-81] | BD #555675 | 0.2 µg per staining | Flow cytometry (beads assay) |
| CD9 [MEM-61] | Sigma #SAB4700092 | 0.2 µg per staining | Flow cytometry (beads assay) |
| CD63 [H5C6] | BD #556019 | 0.2 µg per staining | Flow cytometry (beads assay) |
| Alexa Fluor 488 donkey anti-mouse | Invitrogen #A21202 | 1 µl per staining | Flow cytometry (beads assay) |
| Alexa Fluor 647 donkey anti-mouse | Invitrogen #A31571 | 1 µl per staining | Flow cytometry (beads assay) |
| Alexa Fluor 647 Mouse IgG1, κ Isotype [MOPC-21] | Biolegend #400130 | 0.5 µg per staining | Flow cytometry (beads assay, cells) |
| Alexa Fluor 647 anti-human CD80 [2D10] | Biolegend #305216 | 0.5 µg per staining | Flow cytometry (beads assay, cells) |
| APC Armenian Hamster IgG Isotype [HTK888] | Biolegend #400911 | 0.5 µg per staining | Flow cytometry (beads assay, cells) |
| APC anti-mouse CD80 [16-10A1] | Biolegend #104713 | 0.5 µg per staining | Flow cytometry (beads assay, cells) |
| APC Mouse IgG2b κ Isotype [MPC-11] | BioLegend #400321 | 0.5 µg per staining | Flow cytometry (beads assay, cells) |
| APC anti-human CD274/PD-L1 [29E.2A3] | BioLegend #329707 | 0.5 µg per staining | Flow cytometry (beads assay, cells) |
| APC Rat IgG2bk [RTK4530] | Biolegend #400611 | 0.5 µg per staining | Flow cytometry (beads assay, cells) |
| APC mouse PD-L1 [10F.9G2] | Biolegend #124312 | 0.5 µg per staining | Flow cytometry (beads assay, cells) |
| BV421 Mouse IgG1, κ Isotype Control [X40] | BD #562438 | 0.5 µg per staining | Flow cytometry (beads assay, cells) |
| BV421 Anti-Human OX40 Ligand (CD252) [ik-1] | BD #563766 | 0.5 µg per staining | Flow cytometry (beads assay, cells) |
| Alexa Fluor 647 Rat IgG2b, κ Isotype [RTK4530] | BioLegend #400626 | 0.5 µg per staining | Flow cytometry (beads assay, cells) |
| Alexa Fluor 647 anti-mouse CD252 (OX40L) Antibody [RM134L] | BioLegend #108810 | 0.5 µg per staining | Flow cytometry (beads assay, cells) |
| APC Mouse IgG1, κ Isotype [MOPC-21] | BioLegend #400121 | 0.5 µg per staining | Flow cytometry (beads assay, cells) |
| APC Mouse IgG2b $\kappa$ Isotype [MPC-11] | BioLegend #400321 | 0.5 μg per staining | Flow cytometry (beads assay, cells) |
| APC anti-human CD274/PD-L1 [29E.2A3] | BioLegend #329707 | 0.5 μg per staining | Flow cytometry (beads assay, cells) |
| APC anti-human CD273/PD-L2 [MIH18] | BioLegend #345507 | 0.5 μg per staining | Flow cytometry (beads assay, cells) |
| APC anti-human CD270/HVEM [122] | BioLegend #318807 | 0.5 μg per staining | Flow cytometry (beads assay, cells) |
| APC anti-human Galectin-9 [9M1-3] | BioLegend #348907 | 0.5 μg per staining | Flow cytometry (beads assay, cells) |
| APC anti-human CD80 [2D10] | BioLegend #305219 | 0.5 μg per staining | Flow cytometry (beads assay, cells) |
| APC anti-human CD137L/4-1BBL [5F4] | BioLegend #311505 | 0.5 μg per staining | Flow cytometry (beads assay, cells) |
| APC anti-human CD70 [113-16] | BioLegend #355109 | 0.5 μg per staining | Flow cytometry (beads assay, cells) |
| PE/Cyanine7 Mouse IgG2b $\kappa$ Isotype [MPC-11] | BioLegend #400325 | 0.5 μg per staining | Flow cytometry (beads assay, cells) |
| PE/Cyanine7 anti-human CD275/ICOSL [2D3] | BioLegend #309409 | 0.5 μg per staining | Flow cytometry (beads assay, cells) |
| anti-CD3e [OKT3] | BioLegend #317302 | 1 μg/ml | Functional antibody T cell (ex vivo) |
| anti-CD28 [CD28.2] | eBioscience #16-0289-85 | 1 μg/ml | Functional antibody T cell (ex vivo) |
| anti-CD3e [145-2C11] | BD #553057 | 1 μg/ml | Functional antibody T cell (ex vivo) |
| anti-CD28 [37.51] | BD #553294 | 1 μg/ml | Functional antibody T cell (ex vivo) |
| Fixable Viability Dye eFluor 780 | eBioscience #65-0865-14 | 1 in 1,000 | Flow cytometry |
| APC anti-human CD45 [HI30] | BioLegend #304011 | 1 in 100 | Flow cytometry (ex vivo) |
| PE/Cyanine7 anti-human CD3 [HIT3a] | BioLegend #300316 | 1 in 100 | Flow cytometry (ex vivo) |
| Brilliant Violet 605 anti-human CD4 [RPA-T4] | BioLegend #300556 | 1 in 100 | Flow cytometry (ex vivo) |
| Brilliant Violet 650 anti-human CD8 [RPA-T8] | BioLegend #301042 | 1 in 100 | Flow cytometry (ex vivo) |
| PerCP anti-human CD279 (PD-1) [EH12.2H7] | BioLegend #329937 | 1 in 100 | Flow cytometry (ex vivo) |
| PE/Cyanine7 anti-human CD152 (CTLA-4) [BNI3] | BioLegend #369613 | 1 in 100 | Flow cytometry (ex vivo) |
| APC anti-human CD28 [CD28.2] | BioLegend #302911 | 1 in 100 | Flow cytometry (ex vivo) |
| PE/Dazzle 594 anti-human CD134 (OX40) [Ber-ACT35 (ACT35)] | BioLegend #350020 | 1 in 100 | Flow cytometry (ex vivo) |
| APC-R700 Anti-Human CD25 [2A3] | BD #565106 | 1 in 200 | Flow cytometry (ex vivo) |
| PE-CF594 Anti-Human CD69 [FN50] | BD #562617 | 1 in 200 | Flow cytometry (ex vivo) |
| BV421 Anti-Ki-67 [B56] | BD #562899 | 1 in 100 | Flow cytometry (ex vivo) |
| Brilliant Violet 510 anti-human IL-2 [MQ1-17H12] | BioLegend #500337 | 1 in 100 | Flow cytometry (ex vivo) |
| Brilliant Violet 711 anti-human IFN-γ [4S.B3] | BioLegend #502539 | 1 in 100 | Flow cytometry (ex vivo) |
| FITC anti-human/mouse Granzyme B [GB11] | BioLegend #515403 | 1 in 100 | Flow cytometry (ex vivo) |
| Pacific Blue anti-mouse CD45 [30-F11] | BioLegend #103126 | 1 in 100 | Flow cytometry (ex vivo) |
| Alexa Fluor 700 anti-mouse CD3 [17A2] | eBioscience #56-0032-82 | 1 in 100 | Flow cytometry (ex vivo) |
| Brilliant Violet 605 anti-mouse CD4 [RM4-5] | BioLegend #100548 | 1 in 200 | Flow cytometry (ex vivo) |
| Brilliant Violet 650 anti-mouse CD8 [53-6.7] | BioLegend #100742 | 1 in 200 | Flow cytometry (ex vivo) |
| PerCP/Cyanine5.5 anti-mouse CD279 (PD-1) [29F.1A12] | BioLegend #135208 | 1 in 100 | Flow cytometry (ex vivo) |
| Brilliant Violet 421 anti-mouse CD152 [UC10-4B9] | BioLegend #106312 | 1 in 100 | Flow cytometry (ex vivo) |
| APC anti-mouse CD28 [37.51] | BioLegend #102110 | 1 in 100 | Flow cytometry (ex vivo) |
| PE/Dazzle 594 anti-mouse CD134 (OX-40) [OX-86] | BioLegend #119417 | 1 in 100 | Flow cytometry (ex vivo) |
| APC-R700 Anti-Mouse CD25 [PC61] | BD #565135 | 1 in 200 | Flow cytometry (ex vivo) |
| PE-CF594 Anti-Mouse CD69 [H1.2F3] | BD #562455 | 1 in 200 | Flow cytometry (ex vivo) |
| Brilliant Violet 510 anti-mouse IL-2 [JES6-5H4] | BioLegend #503833 | 1 in 200 | Flow cytometry (ex vivo) |
| BV711 Rat Anti-Mouse IFN-γ [XMG1.2] | BD #564336 | 1 in 200 | Flow cytometry (ex vivo) |
| Pacific Blue anti-mouse CD45 [30-F11] | BioLegend #103126 | 1 in 100 | Flow cytometry (in vivo) |
| Brilliant Violet 510 anti-mouse CD45 [30-F11] | BioLegend #103137 | 1 in 200 | Flow cytometry (in vivo) |
| BV786 Anti-mouse CD11b [M1/70] | BD #740861 | 1 in 200 | Flow cytometry (in vivo) |
| BV711 Anti-mouse CD11b [M1/70] | BD #563168 | 1 in 200 | Flow cytometry (in vivo) |
| Alexa Fluor 700 anti-mouse CD3 [17A2] | eBioscience #56-0032-82 | 1 in 100 | Flow cytometry (in vivo) |
| PE-Cyanine7 anti-mouse CD3e [145-2C11] | eBioscience #25-0031-82 | 1 in 200 | Flow cytometry (in vivo) |
| Brilliant Violet 605 anti-mouse CD4 [RM4-5] | BioLegend #100548 | 1 in 200 | Flow cytometry (in vivo) |
| Brilliant Violet 650 anti-mouse CD8 [53-6.7] | BioLegend #100742 | 1 in 200 | Flow cytometry (in vivo) |
| FITC anti-human/mouse Granzyme B [GB11] | BioLegend #515403 | 1 in 100 | Flow cytometry (in vivo) |
| PE-Cyanine7 T-bet Monoclonal Antibody [4B10] | eBioscience #25-5825-80 | 1 in 200 | Flow cytometry (in vivo) |
| Brilliant Violet 711 anti-mouse CD366 (Tim-3) [RMT3-23] | BioLegend #119727 | 1 in 200 | Flow cytometry (in vivo) |
| APC Rat Anti-Mouse CD62L [MEL-14] | BD #553152 | 1 in 100 | Flow cytometry (in vivo) |
| PE-CF594 Anti-Mouse Foxp3 [MF23] | BD #562466 | 1 in 50 | Flow cytometry (in vivo) |
| PerCP-Cyanine5.5 FOXP3 [FJK-16s] | eBioscience #45-5773-80 | 1 in 50 | Flow cytometry (in vivo) |
| PE-CF594 Hamster Anti-Mouse CD69 [H1.2F3] | BD #562455 | 1 in 200 | Flow cytometry (in vivo) |
| Brilliant Violet 510 anti-mouse IL-2 [JES6-5H4] | BioLegend #503833 | 1 in 200 | Flow cytometry (in vivo) |
| PE anti-mouse Perforin [eBioOMAK-D] | eBioscience #12-9392-80 | 1 in 100 | Flow cytometry (in vivo) |
| BV711 Anti-Mouse IFN-γ [XMG1.2] | BD #564336 | 1 in 200 | Flow cytometry (in vivo) |
| APC-R700 Anti-Mouse CD25 [PC61] | BD #565135 | 1 in 100 | Flow cytometry (in vivo) |
| PerCP-Cy5.5 Anti-Mouse CD19 [1D3] | BD #551001 | 1 in 50 | Flow cytometry (in vivo) |
| Alexa Fluor 488 Mouse anti-Ki-67 [B56] | BD 558616 | 1 in 100 | Flow cytometry (in vivo) |
| InVivoMab Syrian hamster IgG | BioXcell #BE0087 | 200 μg/mouse (D6), 100 μg/mouse (D9, D12, D16) | In vivo treatment |
| InVivoMab anti-mouse CTLA-4 (CD152) [9H10] | BioXcell #BE0131 | 200 μg/mouse (D6), 100 μg/mouse (D9, D12, D16) | In vivo treatment |
| InVivoMab rat IgG2b isotype control [LTF-2] | BioXcell #BE0090 | 200 μg/mouse (D6-D20), 100 μg/mouse (D21-D30) | In vivo treatment |
| InVivoMab anti-mouse CD8α [53-6.7] | BioXcell #BE0004-1 | 200 μg/mouse (D6-D20), 100 μg/mouse (D21-D30) | In vivo treatment |
| InVivoMab rat IgG2a isotype control [2A3] | BioXcell #BE0089 | 200 μg/mouse (D6-D20), 100 μg/mouse (D21-D30) | In vivo treatment |
| InVivoMab anti-mouse CD4 [GK1.5] | BioXcell #BE0003-1 | 200 μg/mouse (D6-D20), 100 μg/mouse (D21-D30) | In vivo treatment |

## References

1 Kalluri, R. & LeBleu, V. S. The biology, function, and biomedical applications of exosomes. Science 367 (2020). 10.1126/science.aau6977

2 Valadi, H. et al. Exosome-mediated transfer of mRNAs and microRNAs is a novel mechanism of genetic exchange between cells. Nat Cell Biol 9, 654–659 (2007). 10.1038/ncb1596

3 Kugeratski, F. G. et al. Quantitative proteomics identifies the core proteome of exosomes with syntenin-1 as the highest abundant protein and a putative universal biomarker. Nat Cell Biol 23, 631–641 (2021). 10.1038/s41556-021-00693-y

4 Zhang, Y., Wu, X. & Andy Tao, W. Characterization and Applications of Extracellular Vesicle Proteome with Post-Translational Modifications. Trends Analyt Chem 107, 21–30 (2018). 10.1016/j.trac.2018.07.014

5 Onozato, M. et al. Amino acid analyses of the exosome-eluted fractions from human serum by HPLC with fluorescence detection. Pract Lab Med 12, e00099 (2018). 10.1016/j.plabm.2018.e00099

6 Haraszti, R. A. et al. High-resolution proteomic and lipidomic analysis of exosomes and microvesicles from different cell sources. J Extracell Vesicles 5, 32570 (2016). 10.3402/jev.v5.32570

7 Ludwig, N., Gillespie, D. G., Reichert, T. E., Jackson, E. K. & Whiteside, T. L. Purine Metabolites in Tumor-Derived Exosomes May Facilitate Immune Escape of Head and Neck Squamous Cell Carcinoma. Cancers (Basel) 12 (2020). 10.3390/cancers12061602

8 Plebanek, M. P. et al. Pre-metastatic cancer exosomes induce immune surveillance by patrolling monocytes at the metastatic niche. Nat Commun 8, 1319 (2017). 10.1038/s41467-017-01433-3

9 Chen, G. et al. Exosomal PD-L1 contributes to immunosuppression and is associated with anti-PD-1 response. Nature 560, 382–386 (2018). 10.1038/s41586-018-0392-8

10 Cooks, T. et al. Mutant p53 cancers reprogram macrophages to tumor supporting macrophages via exosomal miR-1246. Nat Commun 9, 771 (2018). 10.1038/s41467-018-03224-w

11 Gao, L. et al. Tumor-derived exosomes antagonize innate antiviral immunity. Nat Immunol 19, 233–245 (2018). 10.1038/s41590-017-0043-5

12 Ricklefs, F. L. et al. Immune evasion mediated by PD-L1 on glioblastoma-derived extracellular vesicles. Sci Adv 4, eaar2766 (2018). 10.1126/sciadv.aar2766

13 Zhang, F. et al. Specific Decrease in B-Cell-Derived Extracellular Vesicles Enhances Post-Chemotherapeutic CD8(+) T Cell Responses. Immunity 50, 738–750 e737 (2019). 10.1016/j.immuni.2019.01.010

14 Kugeratski, F. G. & Kalluri, R. Exosomes as mediators of immune regulation and immunotherapy in cancer. FEBS J 288, 10–35 (2021). 10.1111/febs.15558

15 Anel, A., Gallego-Lleyda, A., de Miguel, D., Naval, J. & Martinez-Lostao, L. Role of Exosomes in the Regulation of T-cell Mediated Immune Responses and in Autoimmune Disease. Cells 8 (2019). 10.3390/cells8020154

16 Murao, A., Brenner, M., Aziz, M. & Wang, P. Exosomes in Sepsis. Front Immunol 11, 2140 (2020). 10.3389/fimmu.2020.02140

17 Kugeratski, F. G., McAndrews, K. M. & Kalluri, R. Multifunctional Applications of Engineered Extracellular Vesicles in the Treatment of Cancer. Endocrinology 162 (2021). 10.1210/endocr/bqaa250

18 Wang, J., Wang, L., Lin, Z., Tao, L. & Chen, M. More efficient induction of antitumor T cell immunity by exosomes from CD40L gene-modified lung tumor cells. Mol Med Rep 9, 125–131 (2014). 10.3892/mmr.2013.1759

19 Zhang, Y. et al. Exosomes derived from IL-12-anchored renal cancer cells increase induction of specific antitumor response in vitro: a novel vaccine for renal cell carcinoma. Int J Oncol 36, 133–140 (2010).

20 Lewis, N. D. et al. Exosome Surface Display of IL12 Results in Tumor-Retained Pharmacology with Superior Potency and Limited Systemic Exposure Compared with Recombinant IL12. Mol Cancer Ther 20, 523–534 (2021). 10.1158/1535-7163.MCT-20-0484

21 Shi, X. et al. Antitumor efficacy of interferon-gamma-modified exosomal vaccine in prostate cancer. Prostate 80, 811–823 (2020). 10.1002/pros.23996

22 Fu, W. et al. CAR exosomes derived from effector CAR-T cells have potent antitumour effects and low toxicity. Nat Commun 10, 4355 (2019). 10.1038/s41467-019-12321-3

23 Zuo, B. et al. Alarmin-painted exosomes elicit persistent antitumor immunity in large established tumors in mice. Nat Commun 11, 1790 (2020). 10.1038/s41467-020-15569-2

24 McAndrews, K. M., Che, S. P. Y., LeBleu, V. S. & Kalluri, R. Effective delivery of STING agonist using exosomes suppresses tumor growth and enhances anti-tumor immunity. Journal of Biological Chemistry (2021). 10.1016/j.jbc.2021.100523

25 Xing, C. et al. The roles of exosomal immune checkpoint proteins in tumors. Mil Med Res 8, 56 (2021). 10.1186/s40779-021-00350-3

26 Morse, M. A. et al. A phase I study of dexosome immunotherapy in patients with advanced non-small cell lung cancer. J Transl Med 3, 9 (2005). 10.1186/1479-5876-3-9

27 Escudier, B. et al. Vaccination of metastatic melanoma patients with autologous dendritic cell (DC) derived-exosomes: results of thefirst phase I clinical trial. J Transl Med 3, 10 (2005). 10.1186/1479-5876-3-10

28 Zhu, X. et al. Comprehensive toxicity and immunogenicity studies reveal minimal effects in mice following sustained dosing of extracellular vesicles derived from HEK293T cells. J Extracell Vesicles 6, 1324730 (2017). 10.1080/20013078.2017.1324730

29 Mendt, M. et al. Generation and testing of clinical-grade exosomes for pancreatic cancer. JCI Insight 3 (2018). 10.1172/jci.insight.99263

30 Heymann, F., Hamesch, K., Weiskirchen, R. & Tacke, F. The concanavalin A model of acute hepatitis in mice. Lab Anim 49, 12–20 (2015). 10.1177/0023677215572841

31 Larkin, J. et al. Five-Year Survival with Combined Nivolumab and Ipilimumab in Advanced Melanoma. N Engl J Med 381, 1535–1546 (2019). 10.1056/NEJMoa1910836

32 Koh, E. et al. Exosome-SIRPalpha, a CD47 blockade increases cancer cell phagocytosis. Biomaterials 121, 121–129 (2017). 10.1016/j.biomaterials.2017.01.004

33 Yuan, Z., Kolluri, K. K., Gowers, K. H. & Janes, S. M. TRAIL delivery by MSC-derived extracellular vesicles is an effective anticancer therapy. J Extracell Vesicles 6, 1265291 (2017). 10.1080/20013078.2017.1265291

34 Poggio, M. et al. Suppression of Exosomal PD-L1 Induces Systemic Anti-tumor Immunity and Memory. Cell 177, 414–427 e413 (2019). 10.1016/j.cell.2019.02.016

35 Aspeslagh, S. et al. Rationale for anti-OX40 cancer immunotherapy. Eur J Cancer 52, 50–66 (2016). 10.1016/j.ejca.2015.08.021

36 Duhen, R. et al. Neoadjuvant anti-OX40 (MEDI6469) therapy in patients with head and neck squamous cell carcinoma activates and expands antigen-specific tumor-infiltrating T cells. Nat Commun 12, 1047 (2021). 10.1038/s41467-021-21383-1

37 Bansal-Pakala, P., Halteman, B. S., Cheng, M. H. & Croft, M. Costimulation of CD8 T cell responses by OX40. J Immunol 172, 4821–4825 (2004). 10.4049/jimmunol.172.8.4821

38 Sadun, R. E. et al. Fc-mOX40L fusion protein produces complete remission and enhanced survival in 2 murine tumor models. J Immunother 31, 235–245 (2008). 10.1097/CJI.0b013e31816a88e0

39 Gough, M. J. et al. OX40 agonist therapy enhances CD8 infiltration and decreases immune suppression in the tumor. Cancer Res 68, 5206–5215 (2008). 10.1158/0008-5472.CAN-07-6484

40 Kitamura, N. et al. OX40 costimulation can abrogate Foxp3+ regulatory T cell-mediated suppression of antitumor immunity. Int J Cancer 125, 630–638 (2009). 10.1002/ijc.24435

41 Piconese, S., Valzasina, B. & Colombo, M. P. OX40 triggering blocks suppression by regulatory T cells and facilitates tumor rejection. J Exp Med 205, 825–839 (2008). 10.1084/jem.20071341

42 Leach, D. R., Krummel, M. F. & Allison, J. P. Enhancement of antitumor immunity by CTLA-4 blockade. Science 271, 1734–1736 (1996).

43 Iwai, Y., Terawaki, S. & Honjo, T. PD-1 blockade inhibits hematogenous spread of poorly immunogenic tumor cells by enhanced recruitment of effector T cells. Int Immunol 17, 133–144 (2005). 10.1093/intimm/dxh194

44 Ribas, A. & Wolchok, J. D. Cancer immunotherapy using checkpoint blockade. Science 359, 1350–1355 (2018). 10.1126/science.aar4060

45 Wei, S. C., Duffy, C. R. & Allison, J. P. Fundamental Mechanisms of Immune Checkpoint Blockade Therapy. Cancer Discov 8, 1069–1086 (2018). 10.1158/2159-8290.CD-18-0367

46 Varayathu, H., Sarathy, V., Thomas, B. E., Mufti, S. S. & Naik, R. Combination Strategies to Augment Immune Check Point Inhibitors Efficacy - Implications for Translational Research. Front Oncol 11, 559161 (2021). 10.3389/fonc.2021.559161

47 Mascarelli, D. E. et al. Boosting Antitumor Response by Costimulatory Strategies Driven to 4-1BB and OX40 T-cell Receptors. Front Cell Dev Biol 9, 692982 (2021). 10.3389/fcell.2021.692982

48 Shrimali, R. K. et al. Concurrent PD-1 Blockade Negates the Effects of OX40 Agonist Antibody in Combination Immunotherapy through Inducing T-cell Apoptosis. Cancer Immunol Res 5, 755–766 (2017). 10.1158/2326-6066.CIR-17-0292

49 Messenheimer, D. J. et al. Timing of PD-1 Blockade Is Critical to Effective Combination Immunotherapy with Anti-OX40. Clin Cancer Res 23, 6165–6177 (2017). 10.1158/1078-0432.CCR-16-2677

50 Ma, Y. et al. Combination of PD-1 Inhibitor and OX40 Agonist Induces Tumor Rejection and Immune Memory in Mouse Models of Pancreatic Cancer. Gastroenterology 159, 306–319 e312 (2020). 10.1053/j.gastro.2020.03.018

51 Wei, S. C. et al. Distinct Cellular Mechanisms Underlie Anti-CTLA-4 and Anti-PD-1 Checkpoint Blockade. Cell 170, 1120–1133 e1117 (2017). 10.1016/j.cell.2017.07.024

52 Cox, J. & Mann, M. MaxQuant enables high peptide identification rates, individualized p.p.b.-range mass accuracies and proteome-wide protein quantification. Nat Biotechnol 26, 1367–1372 (2008). 10.1038/nbt.1511

53 Cox, J. et al. Andromeda: a peptide search engine integrated into the MaxQuant environment. J Proteome Res 10, 1794–1805 (2011). 10.1021/pr101065j

54 Tyanova, S. et al. The Perseus computational platform for comprehensive analysis of (prote)omics data. Nat Methods 13, 731–740 (2016). 10.1038/nmeth.3901

55 Deutsch, E. W. et al. The ProteomeXchange consortium at 10 years: 2023 update. Nucleic Acids Res 51, D1539–D1548 (2023). 10.1093/nar/gkac1040

56 Perez-Riverol, Y. et al. The PRIDE database resources in 2022: a hub for mass spectrometry-based proteomics evidences. Nucleic Acids Res 50, D543–D552 (2022). 10.1093/nar/gkab1038

